# Knockdown of endothelial *Serpine1* improves stroke recovery by attenuating peri-infarct blood flow and blood brain barrier disruption

**DOI:** 10.1101/2025.10.15.682687

**Authors:** Kamal Narayana, Isabel C. Lambert, Sam Burford, Emilie Gosselin, Jakob Körbelin, Craig E. Brown

## Abstract

Focal stroke leads to complex changes in the cerebral microcirculation in surviving brain tissues that strongly influence recovery. Plasminogen activator inhibitor-1 (PAI-1; encoded by *Serpine1*) is highly upregulated in endothelial cells after stroke. Since the primary function of PAI-1 is to inhibit fibrin clot breakdown, we hypothesized that blocking this pathway would be beneficial for recovery since it is expected to increase capillary blood flow after stroke. Using longitudinal *in vivo* imaging in mice subjected to ischemic stroke, we unexpectedly found that knockdown of *Serpine1* in brain endothelial cells leads to a long-lasting reduction in peri-infarct capillary width, red blood cell velocity and flux. Conversely, stimulating this pathway in naïve mice increased capillary width and blood flow. Lowered peri-infarct blood flow in *Serpine1* knockdown mice attenuated deleterious blood brain barrier disruption and pro-inflammatory gene expression. *Serpine1* knockdown improved the progressive recovery of sensory evoked cortical responses, as well as cognitive and sensorimotor function. These findings challenge the assumption that increased blood flow after stroke is better for recovery and reveal that carefully tuning flow, rather than maximizing it, may be optimal. Further our data highlight the therapeutic potential of targeting endothelial *Serpine1*/PAI-1 signalling in promoting stroke recovery.

## Introduction

Stroke is a cerebrovascular disease that results in irreversible cell death in the ischemic core as well as chronic dysfunction in surviving brain regions^1^. As a result, patients who survive a stroke will likely face permanent disabilities that affect their quality of life. Regulating blood flow in vulnerable brain tissues within the first few hours to weeks after stroke is considered a critical factor in functional recovery^2,3^. However, the relationship between blood flow and functional recovery after ischemic stroke is complex and does not necessarily follow the simple logic that more flow is better. This nuanced relationship is exemplified by the fact that during the hyper-acute phase of ischemic stroke (minutes to hours), restoring some level of blood flow through recanalization of large and small vascular beds or through collaterals, is critical to prevent cell death and long-term functional impairment^4–6^. However, as shown by a recent study from the Wegener group^7^ the restoration of blood flow needs to be carefully tittered, because if it occurs too vigorously, it can lead to micro-hemorrhage and worse stroke outcome in mice and humans. For stroke recovery in the longer term (>24h to several weeks), increased blood flow to surviving peri-infarct regions (above normal levels) may be detrimental, because it could increase the risk of disrupting the blood brain barrier (BBB) and inducing hemorrhage^8^. Indeed, previous work from our lab has shown that diabetic mice, who reliably exhibit increased BBB permeability and worse functional recovery, have persistently elevated blood flow in peri-infarct regions from 3 to 21 days recovery^9^. Collectively these findings suggest that the ideal therapeutic target would provide a sufficient level of blood flow needed to maintain tissue viability and function after stroke, while minimizing complications such as BBB breakdown.

Signaling pathways involved in fibrous clot breakdown such as the *Serpine1* (*Serp1*) gene and its Plasminogen Activator-Inhibitor-1 (PAI-1) protein, are involved in regulating blood flow after stroke. PAI-1 is the primary physiological inhibitor of tissue plasminogen activator (tPA) and urokinase-PA (uPA) in the coagulation cascade. PAI-1 prevents tPA from converting plasminogen to plasmin, effectively preventing the breakdown of fibrinous clots (**Fig. 1A**). PAI-1 levels rapidly rise in blood and brain after stroke^10^, primarily produced from endothelial cells, platelets and to a smaller extent microglia^11–15^. High levels of plasma PAI-1 is a risk factor for ischemic stroke and re-infarction (i.e., secondary stroke) in human patients^16–22^. In addition to regulating coagulation, a much lesser known function of endothelial *Serp1*/PAI-1 is to influence blood vessel tone, at least in heart smooth muscle^23,24^. Thus *Serp1*/PAI-1 signalling is positioned to regulate both coagulation and vessel tone, raising the intriguing possibility that blocking *Serp1*/PAI-1 signalling could be a promising strategy for altering blood flow in a manner that optimizes outcomes after stroke. Furthermore, given its modulatory role in the anti-coagulant pathway, some of the deleterious side effects of thrombolytics may be avoided with inhibiting *Serp1*/PAI-1 signalling.

**Figure 1:**
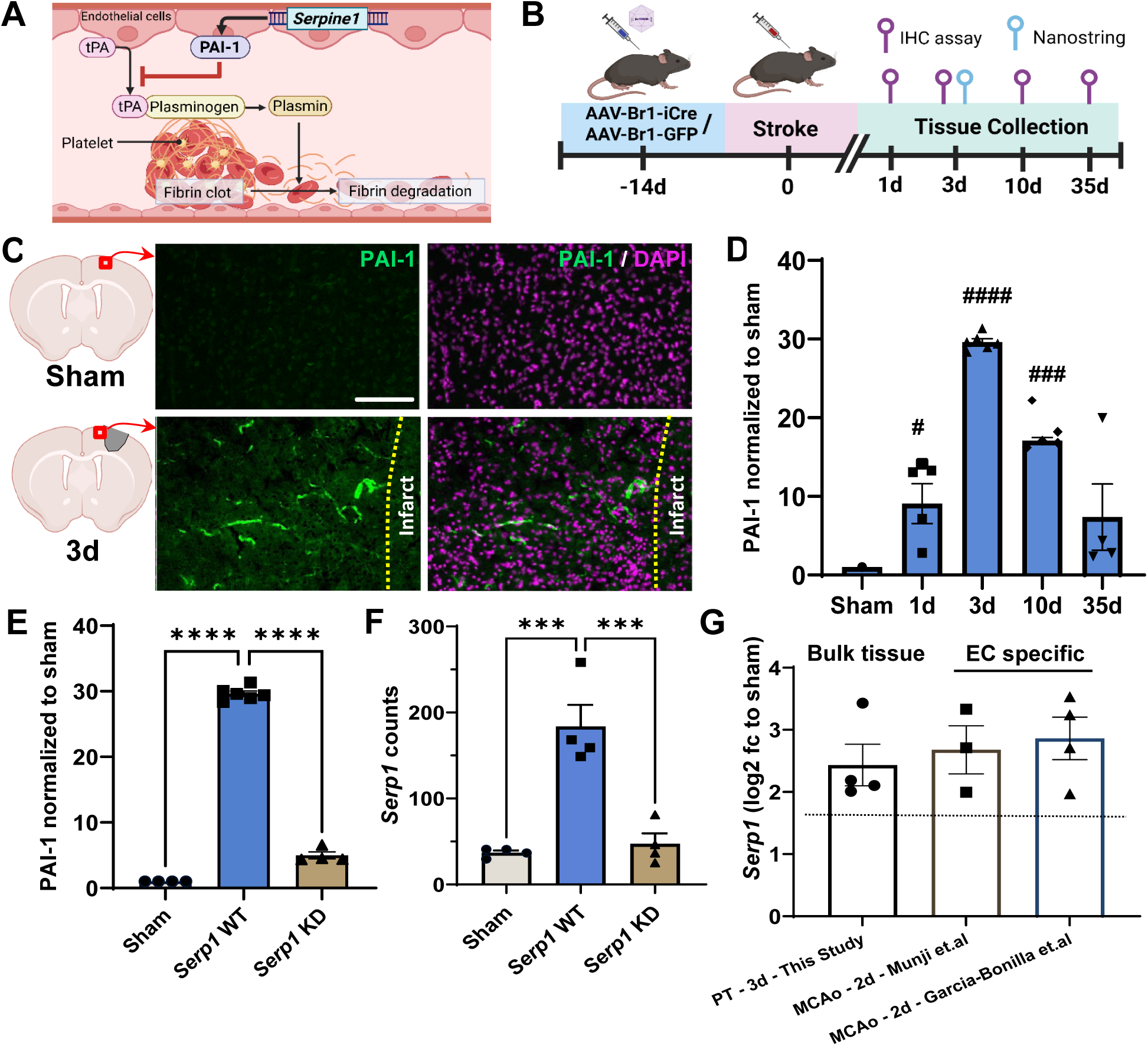
S*e*rpine1/PAI-1 is upregulated in peri-infarct cortex. **A.** Summary diagram showing PAI-I regulation of thrombolysis. **B.** Timeline of experiments examining *Serp1*/PAI-1 changes after stroke. **C.** Confocal immunofluorescent images of PAI-1 (green) and DAPI (purple) in the peri-infarct region (PI; yellow border) in sham stroke control and 3d post-stroke; scale bar = 100µm. **D.** PAI-1 immunostaining area normalized to sham control in peri-infarct regions in sham (*n* = 4), 1d = (*n* = 4), 3d (*n* = 5), 10d (*n* = 4), and 35d (*n* = 4). One-way ANOVA & Sidak’s multiple comparisons comparing post-stroke day to sham; #### *p* < 0.0001, ### *p* < 0.001. **E.** PAI-1 immunostaining area normalized to sham control in peri-infarct cortex of sham (no stroke), *Serp1* WT and KD at 3d post-stroke. One-way ANOVA & Tukey’s multiple comparisons **** *p* < 0.0001. **F.** *Serp1* gene expression levels in peri-infarct cortex of sham (no stroke), *Serp1* WT and KD at 3d post-stroke (*n* = 4/group). One-way ANOVA & Tukey’s multiple comparisons *** *p* < 0.001. **G.** Comparison of *Serp1* gene expression (based on Log2 fold change vs sham controls) at 3d post-stroke (this study) and two separate studies examining endothelial cell (EC) expression at 2d post-stroke. Models of stroke: PT – photothrombotic stroke (this study), MCAo – middle cerebral artery occlusion (Munji et.al (*n* = 3), and Garcia-Bonilla et.al. (*n* = 4)). Data expressed as the mean ± SEM.

There are pre-clinical animal studies that have systemically modulated PAI-1 activity after stroke, but with mixed findings. For example, protective effects of *Serp1*/PAI-1 have been shown given that congenital *Serp1* knockout mice have significantly larger infarcts after focal ischemic stroke while PAI-1 over-expression reduces the extent of infarction^18,25–29^. Contrasting with this, other studies have reported a deleterious role since PAI-I can potentiate neurovascular damage after brain trauma^30^. Further, systemic inhibition of PAI-1 with knockout mice or intravenous injection of blocking antibodies led to improved blood flow in large surface vessels, enhanced behavioural measures of recovery and lowered infarct volumes after ischemic stroke^18,31,32^. However, these studies were limited by the fact that they did not address blood flow or vascular disruption/leakage at the microvascular level, and used non-specific approaches with either a congenital knockout mouse or systemic injection of PAI-I inhibitors. Thus, it remains uncertain whether manipulating *Serp1*/PAI-1 alters microvascular blood flow, influences blood brain barrier (BBB) permeability, or if cell-specific manipulations are effective. In the present study, we focused on inhibiting *Serp1*/PAI-1 signalling specifically within brain endothelial cells given their central role in regulating coagulation and cerebral blood flow. To do this, we longitudinally imaged microvascular networks *in vivo* before and after photothrombotic stroke in mice with viral mediated knockdown of endothelial *Serp1*. Contrary to expectations, we found a long lasting reduction in peri-infarct blood flow in *Serp1* knockdown mice. Reduced blood flow was associated with improved integrity of the BBB and potentiated recovery of cortical hemodynamic responses, sensori-motor and cognitive function.

## Results

### Stroke upregulates *Serpine1* and PAI-1 expression in peri-infarct regions

In order to characterize the temporal expression of *Serp1* mRNA and PAI-1 protein after ischemic stroke, we induced photothrombosis^33^ in the right forelimb somatosensory cortex in 2-4 month old male and female wild-type (WT) mice (**Fig. 1B**). In sham stroke mice, very little PAI-1 protein (**Fig. 1C-E**) or *Serp1* mRNA (**Fig. 1F**) was expressed in cortex. However, ischemic stroke led to a highly significant increase in *Serp1* and PAI-1 expression in peri-infarct cortex that peaked 3 days after stroke (**Fig. 1C-F**). Since our PAI-1 immunostaining indicated a vascular pattern of expression (**Fig. 1C**), but our gene expression assay was based on bulk tissue, we could not definitively conclude that *Serp1* was upregulated in endothelial cells. Therefore, we examined *Serp1* gene expression in published single cell RNAseq datasets collected after ischemic stroke^11,12^. These databases confirm our finding showing that *Serp1* is significantly upregulated in peri-infarct vascular endothelial cells at 2 days after permanent middle cerebral artery occlusion (**Fig. 1G**).

Given that stroke increases *Serp1*/PAI-1 expression in peri-infarct endothelial cells, we next focused on endothelial specific manipulations of *Serp1*. To knock down *Serp1* gene expression in brain endothelial cells, *Serp1* floxed mice were injected (i.v.) with AAV-BR1-iCre (“*Serp1* KD”) or control virus (AAV-BR1-eGFP, “*Serp1* WT”) two weeks before the induction of stroke (**Fig. 1B**). Previous research has shown that AAV-BR1 infects endothelial cells in the brain but not other cell types like microglia or other organs^34^. This cellular specificity in viral infection is important since *Serp1* is also expressed in microglia. Examination of peri-infarct tissues at 3 days post-stroke showed that PAI-1 and *Serp1* expression were significantly decreased in mice with *Serp1* KD mice compared to WT controls (**Fig. 1E,F**). Furthermore, we functionally validated the knockdown of *Serp1* in cerebral capillaries by examining re-flow in capillaries that had clotting induced by targeted 2-photon irradiation (**Supp. Fig. 1**). We reasoned this was a more relevant assay than measuring clotting time in the conventional tail bleed tests. In wild-type mice, 100% of capillaries remained clotted at 6 hours (10/10 capillaries in 2 mice) as shown by the absence of blood flow. By contrast only 41.7% of capillaries in *Serp1* KD mice remained clotted (10/24 capillaries in 2 mice; ꭓ^2^(1,34) = 7.66, p=0.006). Collectively, these experiments validate AAV-mediated *Serp1* knockdown by showing: a) *Serp1* and PAI-1 expression after stroke is significantly reduced and b) clotted cerebral capillaries are more likely to recanalize in *Serp1* KD mice, thus providing functional evidence of reduced PAI-1 activity.

### Endothelial *Serpine1* knockdown reduces capillary blood flow in peri-infarct cortex

Following validation of *Serp1* KD, we next investigated the role endothelial *Serp1*/PAI-1 signalling plays in stroke damage and recovery. First we quantified the volume of ischemic damage at 3 days post stroke. Our analysis showed that infarct volume in *Serp1* KD mice was not significantly different than in *Serp1* WT mice (**Fig. 2A**). Since we did not observe any acute neuroprotective effects of *Serp1* KD, we next focused on microvascular blood flow dynamics that evolve over days and weeks after stroke^35^. To do this, we implanted cranial windows in homozygous *Serp1* floxed mice and then injected them with AAV-BR1-iCre or GFP to yield *Serp1* KD or *Serp1* WT mice, respectively (**Fig. 2B**). Under light isoflurane anesthesia, blood flow dynamics in microvascular cortical networks were imaged before the induction of photothrombotic stroke in forelimb somatosensory cortex and then re-imaged 3, 10, 21 and 35 days afterwards (**Fig. 2B-D**). The sham stroke control group consisted of both *Serp1* WT and KD mice (n=6 total with 3 mice from each genotype) since *Serp1*/PAI-1 expression is extremely low in the absence of stroke and RBC velocities did not differ between these two types of controls over time (**Supp** **Fig. 2**). We focused our imaging on peri-infarct regions that were approximately 184 ± 21µm from the infarct border (range from 113 to 352µm), as well as more distant regions located 849 ± 58µm (range from 491 to 1643µm). We first quantified RBC velocity (mm/s) across all capillaries using line scans, where the streaking pattern of unlabelled RBCs moving through the capillary lumen can be used to estimate RBC velocity (**Fig. 2C**). As shown in Figure 2E, RBC velocity before stroke (“Pre-stroke”) was similar across the three experimental groups (F(4, 1152) = 1.98, *p* = 0.28). In mice subjected to sham stroke procedure, RBC velocities did not change significantly over the 35 day imaging period (**Fig. 2E** **and Supp Fig. 2**; Note that since there is no peri-infarct or distant region in sham controls, the same data is plotted in both graphs).

**Figure 2:**
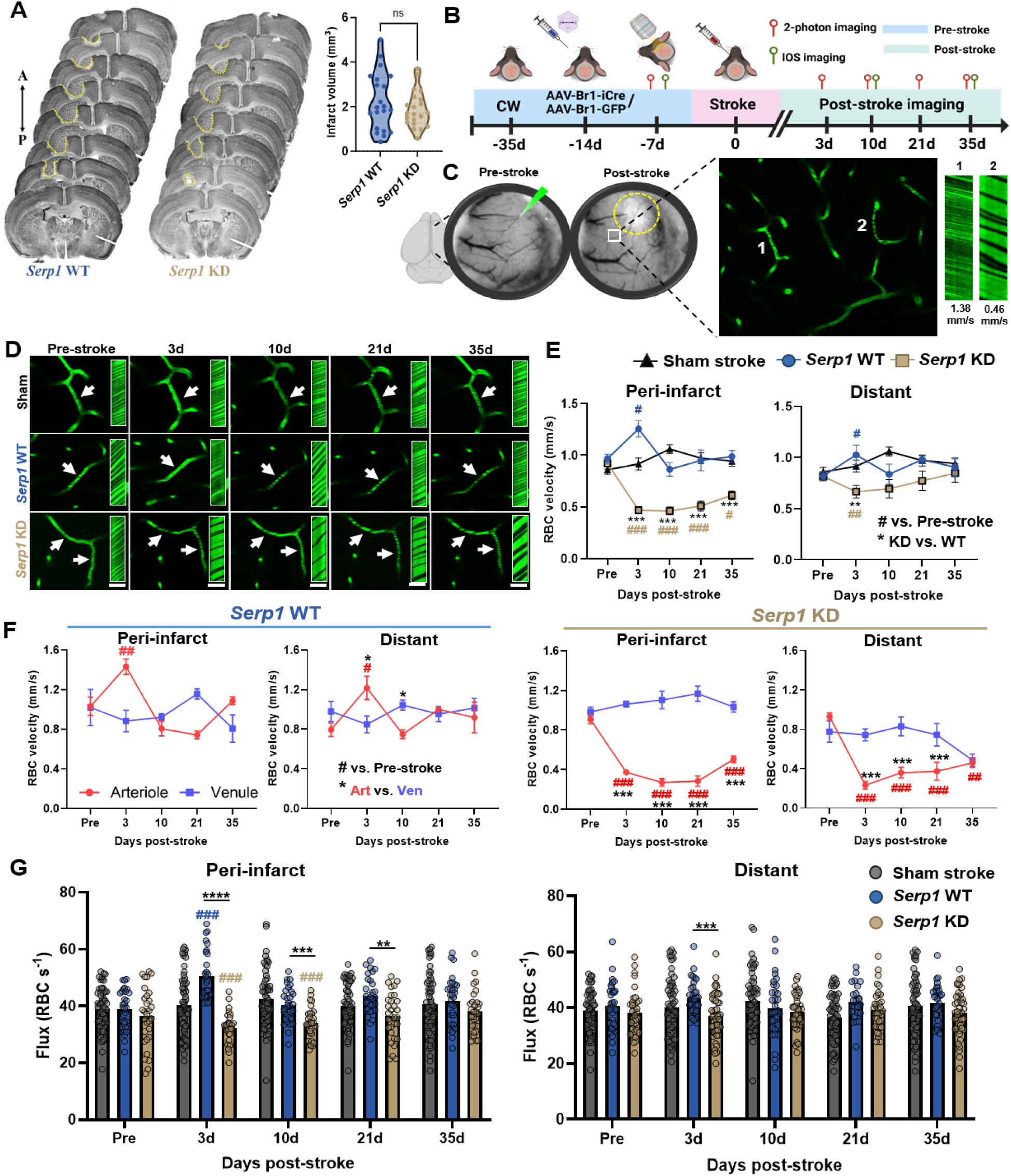
S*e*rpine1 KD leads to reduced blood flow velocity and flux in peri-infarct regions. **A.** Infarcts (denoted by yellow border) were measured in *Serp1* WT (blue) and *Serp1* KD (golden) mice 3 days after stroke (n = 7/group). Data analyzed with two-tailed independent *t*-test; ns, *p* = 0.51. **B.** Timeline of longitudinal *in vivo* imaging experiments. **C.** Brightfield image showing brain surface before and 3 days after stroke. Boxed region shows a maximum intensity z-projection of 40 planar images (1.25 µm apart) alongside with line scans from 2 capillaries; one fast flowing (1.38mm/s), and the other slow (0.46mm/s). Scale bar = 50µm. **D.** *In vivo* maximum z-projection images illustrating blood flow changes across the 35-day imaging period. Scale bar = 50µm. **E.** RBC velocity measurements in capillaries from sham control or peri-infarct and distant regions across 35 day stroke recovery period. N = 6 sham stroke mice, N = 10 and 17 stroke affected *Serp1* WT and KD mice, respectively. **F.** RBC velocity measurements in stroke affected *Serp1* WT and KD capillaries binned by proximity to nearest penetrating arteriole (red lines) or ascending venule (purple lines). **G.** RBC flux (RBCs/s) in capillaries from sham stroke controls or stroke affected *Serp1* WT (blue) and KD mice (golden) over time. 81 capillaries were measured in each group. Data in **E-G** were analysed with two-way ANOVA followed by Tukey’s multiple comparisons test. Comparisons between *Serp1* KD vs WT: **** *p* < 0.0001, *** *p* < 0.001, ** *p* < 0.01, * *p* < 0.05. Comparisons between post-stroke time-point to their respective pre-stroke (0 day) value: #### *p* < 0.0001, ### *p* < 0.001, ## *p* < 0.01, # *p* < 0.05. Data expressed as the mean ± SEM.

However, RBC velocities showed strikingly different patterns after stroke when comparing *Serp1* WT to KD (**Fig. 2E**). These velocity differences between genotypes were most apparent for peri-infarct regions (2-way ANOVA, Main effect of Genotype: F(1, 638) = 130.9, *p* < 0.0001; Main effect of Time: : F(4, 638) = 9.97 *p* < 0.0001; Genotype x Time Interaction: F(4, 638) = 13.25, *p* < 0.0001) and to a lesser extent in distant regions (2-way ANOVA, Main effect of Genotype: F(1, 286) = 7.79, *p* = 0.006; No significant effects of Time or Genotype x Time interaction). In wild-type mice, RBC velocities increased significantly at 3 days in peri-infarct and distant regions (see blue hashtags in **Fig. 2E**), but returned to baseline levels for the remaining post-stroke imaging sessions. By contrast in *Serp1* KD mice, RBC velocities decreased significantly at 3 days and remained well below pre-stroke values for the entire imaging period in the peri-infarct region (see gold hashtags in **Fig. 2E**). Given well known differences in contractility depending on whether capillaries branch off a penetrating arteriole (PA) verses an ascending venule^36,37^ (AV) we grouped RBC velocity measurements in capillaries based on whether they branched off the nearest PA or AV. Before stroke induction, RBC velocities were similar in capillary branches proximal to PA or AVs, regardless of genotype or where they were measured form (**Fig. 2F**).

Following stroke, the biggest changes in RBC velocity originated from capillaries proximal to the PA in both peri-infarct and distant regions (**Fig. 2F**). Not surprisingly, the most significant differences occurred within *Serp1* KD mice (right panel in **Fig. 2F**). And finally, we examined RBC flux (# RBC/s) in capillaries before and after stroke (**Fig. 2G**). Consistent with the observed reduction in RBC velocity, *Serp1* KD mice showed significantly lower RBC flux at 3, 10 and 21 days in peri-infarct regions relative to *Serp1* WT mice (**Fig. 2G**). To eliminate the possibility that AAV-BR1-iCre viral infection could independently affect blood flow, we confirmed that the RBC velocity did not fluctuate in *Serp1* KD without stroke (sham) across the 35 days (avg. velocity = 0.93 mm/s, p = 0.78) (**Supp. Fig. 2**). These results demonstrate endothelial knockdown of *Serp1* causes a significant reduction in RBC velocity and flux in peri-infarct cortex.

### Endothelial *Serpine1* knockdown induces prolonged capillary constriction after stroke

To better understand if reduced velocity/flux was related to constriction/narrowing of capillaries, we repeatedly measured widths from the first capillary branch of a PA or AV, down to the sixth order branch, both before and after induction of stroke (**Fig. 3A,B**). As expected from previous work^38^, capillary widths decreased with increasing branch orders, going from an average diameter of ∼7 to ∼3 µm (**Fig. 3C,D**). Consistent with our RBC velocity and flux measurements in *Serp1* WT, as well as previous imaging work^9^, we noted a transient dilation of 1^st^-3^rd^ order capillaries 3 days after stroke in WT mice (see blue hashtags at day 3 in **Fig. 3C**). Conversely, *Serp1* KD mice showed a highly significant and persistent reduction in capillary widths after stroke that was most evident in branch orders arising from the PA (top row in **Fig. 3C**), with less robust changes in branches off the AV (bottom row in **Fig. 3C**). When examining capillary widths in more distant stroke regions (**Fig. 3D**), similar trends were observed although the magnitude of stroke and genotype based differences were generally reduced. To confirm the potential dilatory effects of PAI-1, we allowed for recombinant PAI-1 (rPAI-1) to diffuse in *Serp1* WT mice (ie. baseline levels of PAI-1) throughout the brain and followed this up *in vivo* measuring vessel width and blood flow 6 and 24hrs post-treatment (**Fig. 3E-F**). We measured a significant increase in vessel width at 6 hrs post-treatment but not at 24hrs (1-way ANOVA, Main effect of Time to Treatment: F(2, 121) = 3.355, *p* = 0.038) (**Fig. 3G**), and when normalized to pre-treatment (−7 days), there was a 9.9% increase in vessel width at 6 hrs only (Left graph shows individual vessel; paired t-test, Right graph shows mean; one sample t-test, *t* = 3.222, df = 55, *p* = 0.0025) (**Fig. 3H**). In addition, blood flow also significantly increased at 6 hrs only (1-way ANOVA, Main effect of Time to Treatment: F(2, 30) = 4.256, *p* = 0.0236) (**Fig. 3I**). To conclude, *Serp1* KD mice exhibit an unusual stroke related constriction of capillaries, particularly in those arising from the PA in peri-infarct cortex. Moreover, diffusion of rPAI-1 lead to a moderate increase in vessel width 6 hrs post-treatment validating the dilatory effects of PAI-1.

**Figure 3:**
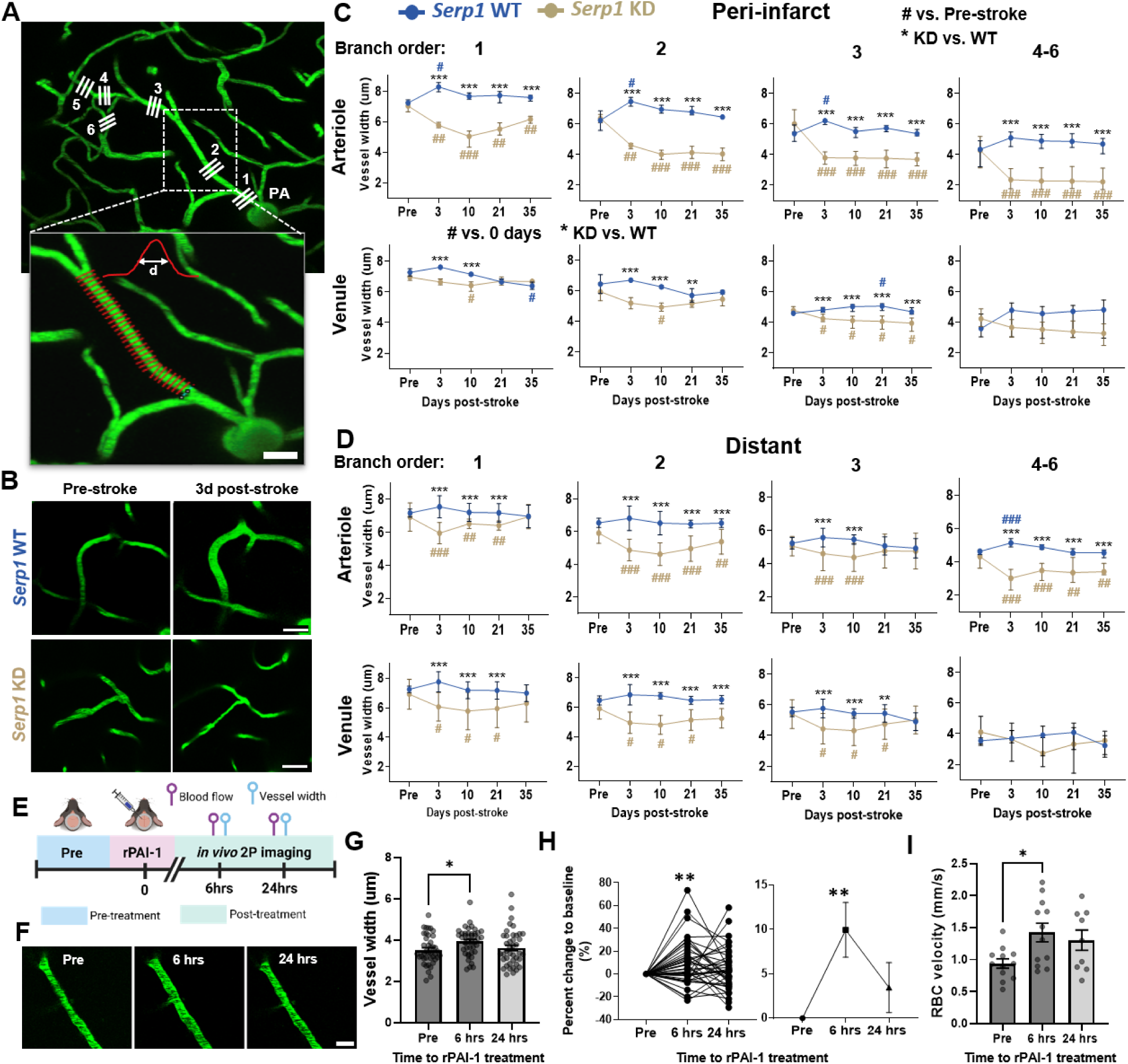
Endothelial *Serpine1* KD leads to prolonged constriction in peri-infarct capillaries. **A.** *In vivo* maximum intensity z-projection images showing a penetrating arteriole (PA) and downstream branches from 1^st^ to 6^th^ order. Boxed region shows example of capillary diameter (d) measurements using Vasometrics tool in ImageJ/Fiji where diameter is estimated based on half-maximal intensity from linear plots of intravascular fluorescence. Scale bar = 50µm. **B.** Representative maximum intensity z-projection images showing that peri-infarct capillaries dilate 3 days after stroke in *Serp1* WT mice but constrict in *Serp1* KD mice. Scale bar = 50µm. **C.** Graphs show capillary widths in peri-infarct cortex of *Serp1* WT (blue) and KD (golden) mice based on their branch order and whether they arise from the PA or AV. **D.** Capillary widths in distant cortex of *Serp1* WT (blue) and KD (golden) mice based on their branch order and whether they arise from the PA or AV. **E.** Timeline of the rPAI-1 *in vivo* experiment. **F**. Example z-projection image of a vessel dilating post-rPAI-1 treatment. Scale bar = 25µm. **G.** Graph showing vessel width in *Serp1* WT mice across the imaging time points. **H**. Graphs showing percent change of vessel width at 6 and 24 hrs when normalized to baseline (−7 days, pre-treatment; %) of individual vessels (left) and mean (right). **I.** RBC velocity measurements in capillaries across the imaging time points. Data in **C-D** were analysed with two-way ANOVA followed by Tukey’s multiple comparisons test. Comparisons between *Serp1* KD vs WT: **** *p* < 0.0001, *** *p* < 0.001, ** *p* < 0.01, * *p* < 0.05. Comparisons between post-stroke time-point to their respective pre-stroke value: #### *p* < 0.0001, ### *p* < 0.001, ## *p* < 0.01, # *p* < 0.05. Data expressed as the mean ± SEM. N = 6 sham stroke mice, N = 10 and 17 stroke affected *Serp1* WT and KD mice, respectively. Data in **G and I** were analysed with one-way ANOVA followed by Tukey’s multiple comparisons test. Comparisons between time of rPAI-1 treatment: * *p* < 0.05. Data in **H** was analysed with a paired t-test comparing to pre-treatment, and one-sample t-test compared to pre-treatment as a hypothetical of zero. Comparisons between time of rPAI-1 treatment: ** *p* < 0.01. Data expressed as the mean ± SEM. N = 4 *Serp1* WT mice, n = 56 vessels for **G-H**, and n = 12 vessels for **I**.

### Endothelial *Serpine1* knockdown reduces BBB permeability after stroke

A reduction in peri-infarct blood flow velocity and flux after stroke could be beneficial for recovery if it were to reduce BBB breakdown, which is well known to have deleterious effects on local synaptic circuits and stroke recovery^39^. To assess permeability of the BBB *in vivo*, we quantified the time dependent extravasation of FITC-dextran (4 kDa) labelled blood plasma in peri-infarct cortex at different time points after stroke^40,41^. As shown in Figure 4A and B, FITC dextran dye slowly accumulates in extravascular space over the 30min imaging period. At 3 days post stroke, BBB permeability was significantly elevated in both *Serp1* WT and KD mice relative to sham stroke controls (**Fig. 4B**). However, BBB permeability was significantly reduced in *Serp1* KD mice compared to WT mice (**Fig. 4B**). Imaging at later recovery time points (10 and 21 days) did not reveal any differences based on genotype (**Fig. 4B**). However, it was apparent that plasma leakage at 10 and 21 days had greatly diminished and approximated permeability levels found in sham control mice, consistent with previous data showing that the BBB re-seals over time^39^. And finally, it is important to note that we did not find evidence of larger scale vascular disruption in the form of hemorrhages in peri-infarct cortex after *Serp1* knockdown.

**Figure 4:**
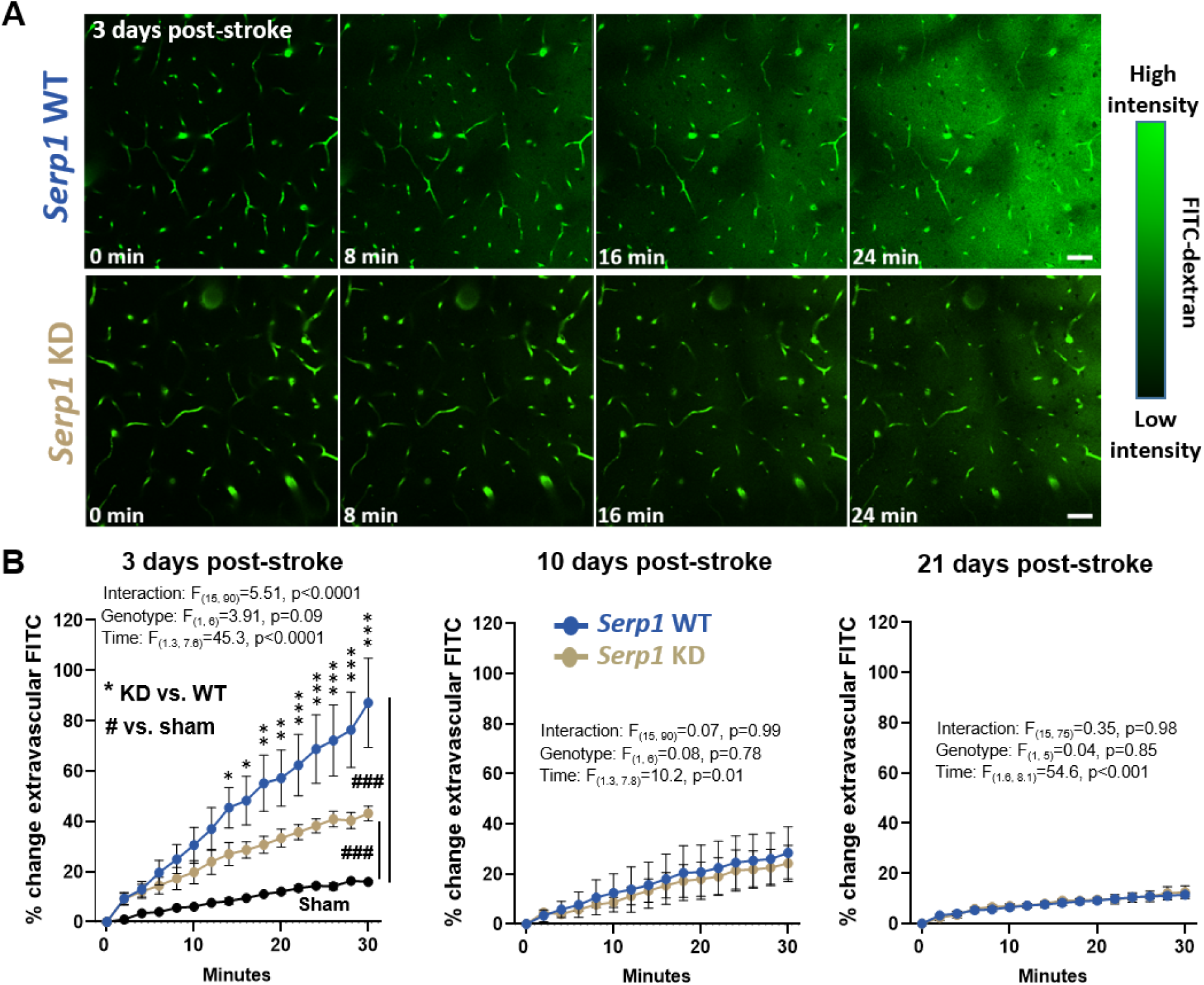
S*e*rpine1 KD reduces blood-brain barrier permeability in peri-infarct cortex. **A.** *In vivo* maximum intensity z-projection images showing FITC-dextran extravasation in peri-infarct cortex of *Serp1* WT and KD mice at 3 days post-stroke (n=6 mice per group). Scale bar = 50µm. **B.** Graphs show the percent change in extravascular FITC in *Serp1* WT and KD mice (n=6 mice/group) at 3, 10 and 21 days post-stroke. Sham stroke controls are plotted in black for comparison (n=4 mice based on 2 WT and 2 KD). Data were analysed with two-way ANOVA followed by Tukey’s multiple comparisons test. Comparisons between *Serp1* KD vs WT: *** *p* < 0.001, ** *p* < 0.01, * *p* < 0.05. Comparisons to sham stroke controls: $$$ *p* < 0.001. Data expressed as the mean ± SEM.

### Endothelial *Serpine1* knockdown alters inflammation related gene and protein expression

Given that endothelial knockdown of *Serp1* significantly reduced BBB permeability at 3 days recovery, we next asked whether this would be associated with a change in inflammation related gene expression in peri-infarct cortex or blood cytokine levels. First, we characterized differentially expressed genes (DEGs) in *Serp1* WT and *Serp1* KD in peri-infarct tissue at 3 days post-stroke (n=4 mice/group, see **Fig. 2B** for experimental summary) and compared them to sham stroke controls consisting of both *Serp1* WT and KD mice (n=4 total with 2 mice from each genotype). As expected from previous studies examining transcriptomic changes in cortical tissue after stroke, several injury related genes were significantly upregulated in *Serp1* WT mice after stroke (**Supp** **Fig. 3**), including *Serp1* (in WT only)*, Serpina3n, Spp1 Cd68, Cd74, Slamf9, Lcn2, Ccl2, Mmp12* and *Timp1*. *Serp1* KD mice displayed a slightly different expression profile (**Supp** **Fig. 3**) although some gene expression changes were common to both genotypes (Upregulated: *Serpina3n*, *Cd68, Slamf9*; Downregulated: *Shank3, Vegfa, Nwd1, Pink1*). Directly comparing gene expression between *Serp1* WT and KD mice at 3 days post-stroke indicated that many of these injury and inflammation related genes (including *Serp1*) were significantly decreased in *Serp1* KD mice relative to WT (**Fig. 5A-C**). Gene ontology analysis supported the finding that inflammation related biological processes and molecular functions were reduced in *Serp1* KD mice (**Fig. 5D**). Similarly, KEGG analysis showed that gene functions most strongly down-regulated in *Serp1* KD mice were associated with microglia function, innate and adaptive immune responses, as well as Notch and growth factor signalling (**Fig. 5E**). In addition to genomic alterations, we also analyzed cytokine levels in blood serum collected 3 days after stroke or sham stroke procedure. As shown in Figure 5F, stroke led to a significant increase in IFN-γ, M-CSF, IL-6, and IL-12p40 in *Serp1* WT, whereas only IFN-γ, IL-2, IL-10, and IL-12p40/70 were elevated in *Serp1* KD mice. When examining differences between genotypes (see asterisks in **Fig. 5F**), *Serp1* KD mice had significantly lower levels of M-CSF, IL-6 and IL-12p40 relative to *Serp1* WT mice, whereas IL-2, IL-10, and IL-12p70 were significantly higher in *Serp1* KD mice. Collectively these results suggests that *Serp1* knockdown leads to a reduction in inflammation related gene expression in peri-infarct cortex and lower levels of putative pro-inflammatory cytokines in blood plasma.

**Figure 5.**
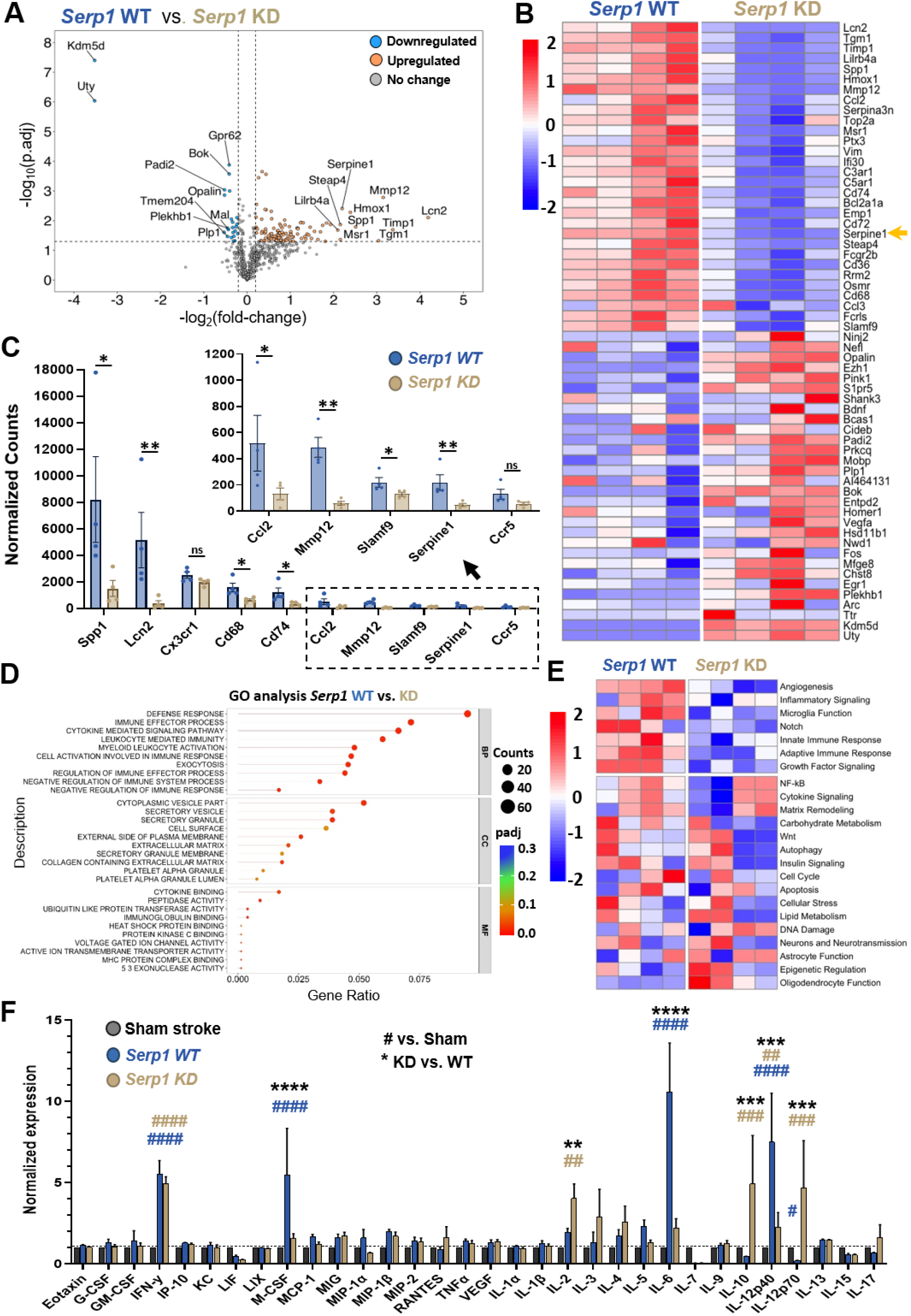
*Serpine1* KD alters inflammatory gene and cytokine expression after stroke. **A.** Volcano plot of 770 differentially expressed genes (DEGs) in peri-infarct tissue of *Serp1* WT versus KD normalized to sham stroke (n=4 mice/group), with the top 10 most up-(orange) and down-regulated (blue) labelled. Data is analysed with moderated t-test with multiple test corrections using limma with false discovery rate (FDR)<0.05 and an absolute value of log2 FC > 0.3 is colored as dotted lines. **B.** Heatmap showing log2 fold change (FC) of the top-30 most significantly up- and down-regulated genes in both genotypes (WT-left, KD-right). Scale for relative gene expression changes range from +2 (up-regulated, red) to -2 (down-regulated, blue). **C.** Histogram shows normalized gene counts of the top-10 most significantly regulated inflammatory genes (*padj* < 0.05 for all comparisons between genotypes). Data expressed as the mean ± SEM. **D.** Dot plot showing GO enrichment analysis of DEGs (two-sided test) for the most significantly regulated processes in *Serp1* WT vs KD; BP – biological process, CC – cellular component, and MF – molecular function. **E.** Heatmap of log2 fold change of the most regulated pathways in both genotypes. The top 7 most differentially regulated processes/pathways between groups is shown in top panel. DEGs in **B** and **E** with FDR<0.05 were ranked by their log2FC, and z-scores were computed on average gene expression for visualization in heatmaps. **F.** Cytokine expression in blood serum at 3d post-stroke, normalized to sham control (n=4 mice/group). Data in F analysed with two-way ANOVA and Tukey multiple comparisons. Comparisons between *Serp1* KD vs WT: **** *p* < 0.0001, *** *p* < 0.001, ** *p* < 0.01. Comparisons between each genotype and sham stroke control: #### *p* < 0.0001, ### *p* < 0.001, ## *p* < 0.01.

### Endothelial *Serpine1* knockdown promotes recovery of somatosensory cortical function

Based on the evidence that *Serp1* KD acutely (i.e. at 3 days recovery) reduced BBB permeability and inflammation related gene and protein expression after stroke, we next probed functional indicators of stroke recovery in the longer term (see summary diagram in **Fig. 1B**). Previous studies have shown that the recovery of cortical responses and activity patterns in the stroke affected hemisphere are predictive of behavioural recovery^42–44^. To this end, we imaged sensory-evoked intrinsic optical signals (IOS) in *Serp1* WT and KD mice before stroke and then again at 10 and 35 days recovery. IOS signals (at least the initial dip in signal) reflect local increases in deoxyhemoglobin levels due to an increase in sensory evoked neuronal activity, thus providing an estimate of functional cortical activity/responses after stroke. However since IOS signals can be influenced by systemic changes in cardiovascular function, we first determined that there were no significant differences between genotypes in oxygen saturation, heart rate and breath rate (**Supp** **Fig. 4**). We also analysed blood chemistry, in particular hemoglobin and hematocrit which can influence light absorbance in IOS imaging. This analysis did not reveal any genotypic differences in hematocrit and hemoglobin (**Supp Fig. 5**), although there were some subtle group differences in Na^+^ and HCO_3_ levels which were still within the normal range. In the absence of stroke, vibrotactile stimulation of the fore- or hindlimb (FL and HL, respectively) elicits a robust and focal response in primary somatosensory cortex in mice before stroke (**Fig. 6A**). The amplitude of sensory evoked responses before stroke did not differ between genotypes for both FL (F(1, 2617)=13.21, *p* > 0.99) and HL cortex (F(1, 4352)=10.54, *p* > 0.99). At 10 days post-stroke, FL and HL cortical responses were significantly reduced in amplitude in both genotypes (**Fig. 6B,C**). However the loss of cortical responses at 10 days was significantly blunted in *Serp1* KD mice relative to *Serp1* WT (**Fig. 6B,C**). By 35 days of recovery, the amplitude of *Serp1* KD IOS responses had recovered back to pre-stroke levels (unpaired t-test compared pre-stroke to day 35 in FL and HL: *p* = 0.64 and 0.89, respectively) whereas *Serp1* WT responses still remained significantly below baseline levels (see blue hashtags in **Fig. 6C**). These results indicate that *Serp1* KD helps preserve and/or recover sensory cortical function after stroke.

**Figure 6:**
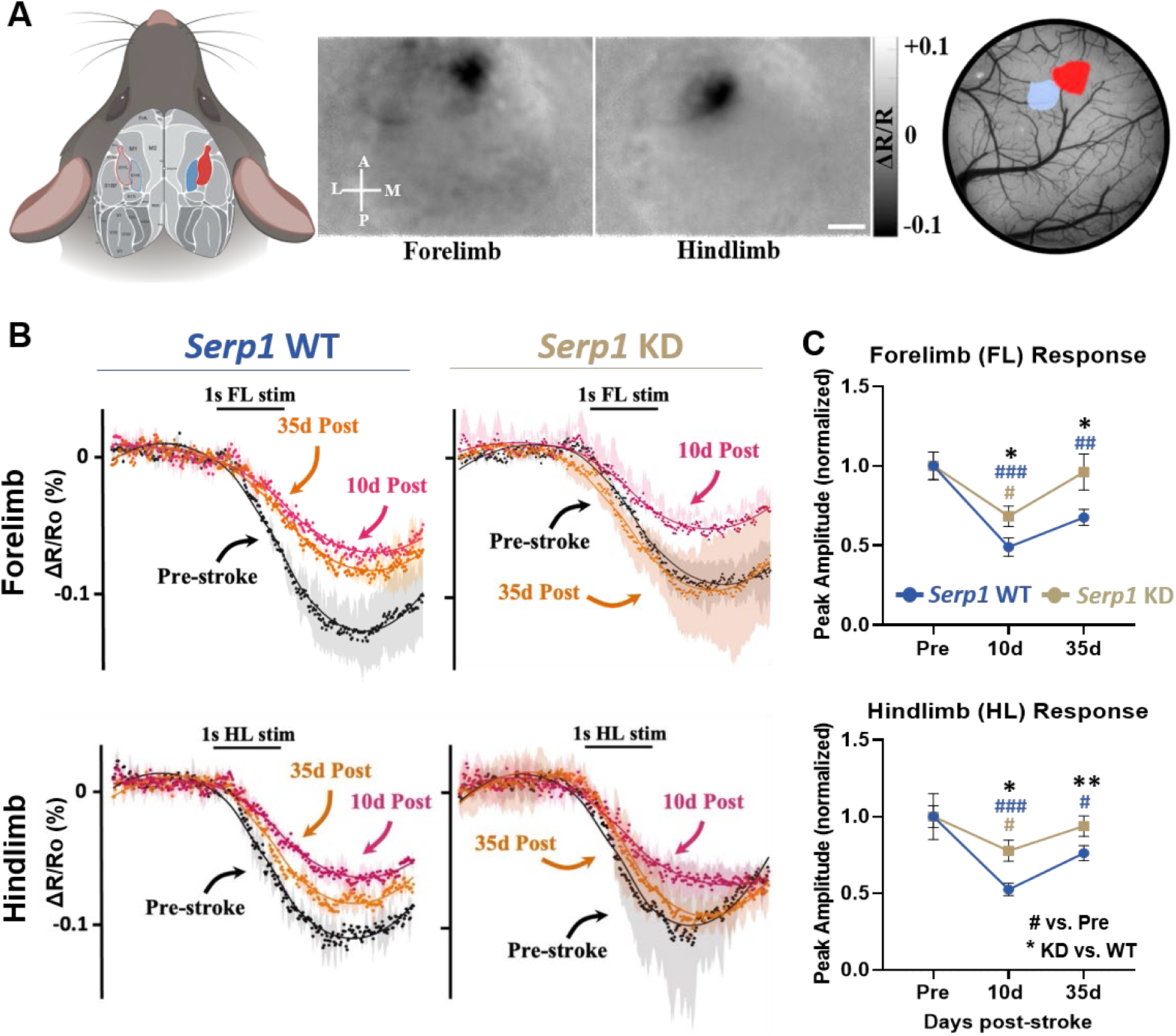
S*e*rpine1 KD mice show enhanced recovery of sensory evoked cortical responses after stroke. **A.** Left: Mouse brain shows a map of forelimb (FL, red) and hindlimb (HL, blue) regions. Middle: representative FL and HL evoked IOS signals based on changes in light reflectance relative to pre-stimulus baseline (% ΔR/Ro). Each response was thresholded and super-imposed onto the cortical surface image. Scale bar = 1mm. **B.** Graphs show average change (% ΔR/Ro) of the FL and HL evoked cortical response in *Serp1* WT (left, blue, n=7 mice) and *Serp1* KD (right, golden, n=10 mice) measured pre-stroke (black), 10d (pink) and 35d (orange) post-stroke. **C.** Graphs show peak amplitude of FL and HL evoked responses in each genotype normalized to pre-stroke values at 10d and 35d after stroke. Data in **C** analyzed with two-way ANOVA and Tukey’s multiple comparisons test. Comparisons between *Serp1* KD vs WT: ** *p* < 0.01, * *p* < 0.05. Comparisons between each genotype and Pre-stroke value: ### *p* < 0.001, ## *p* < 0.01, #p<0.05. Data expressed as the mean ± SEM.

### *Serpine1* knockdown improves behavioural recovery from ischemic stroke

Next we subjected mice to a battery of sensori-motor and cognitive behavioural tests to determine if *Serp1* KD ultimately improved function after stroke (**Fig. 7A**). First, we examined forepaw function using the adhesive tape removal and horizontal ladder walking tests (**Fig. 7B,C**). For the tape removal test, stroke led to a significant increase in tape removal time for both genotypes (**Fig. 7B**). However, *Serp1* KD mice exhibited significantly reduced tape removal latencies from day 10 onwards (from the stroke affected left paw) compared to *Serp1* WT mice (**Fig. 7B**). In the horizontal ladder test, stroke led to a significant increase in the percentage of partial/incorrect forepaw placements in both *Serp1* WT and KD mice when compared to sham controls (**Fig. 7C**). At later recovery time points, *Serp1* KD exhibited marginally better performance on this test relative to WT mice, which was significant at 35 days (**Fig. 7C**). To assess spatial learning and memory, we tested mice in the Morris water maze (MWM) at 7 and 35 days post-stroke (**Fig. 7D**). When examining how long it took mice to find the submerged platform, *Serp1* KD mice performed significantly better than *Serp1* WT mice at both 7 days (**Fig. 7D**; 2-way ANOVA, Genotype: F(1, 51)=10.12, *p* =0.002) and 35 days recovery (2-way ANOVA, Genotype: F(1, 49)=6.48, *p* =0.01). In order to assess cognitive flexibility, we switched the submerged platform to a new location in the MWM test at day 37 (ie. “reversal learning”). *Serp1* KD mice took significantly less time to find the (newly located) platform compared to *Serp1* WT mice (**Fig. 7E**, 2-way ANOVA, Genotype: F(1, 50)=5.66, *p* =0.02). And finally, to determine if there were any general changes in locomotor activity and exploration, we performed the open field test at 7d and 35d post-stroke. This test did not reveal any differences based on genotype (**Supp** **Fig. 6**). In summary, our data shows that endothelial knockdown of *Serp1* generally improves metrics of sensori-motor and cognitive function in stroke affected mice.

**Figure 7:**
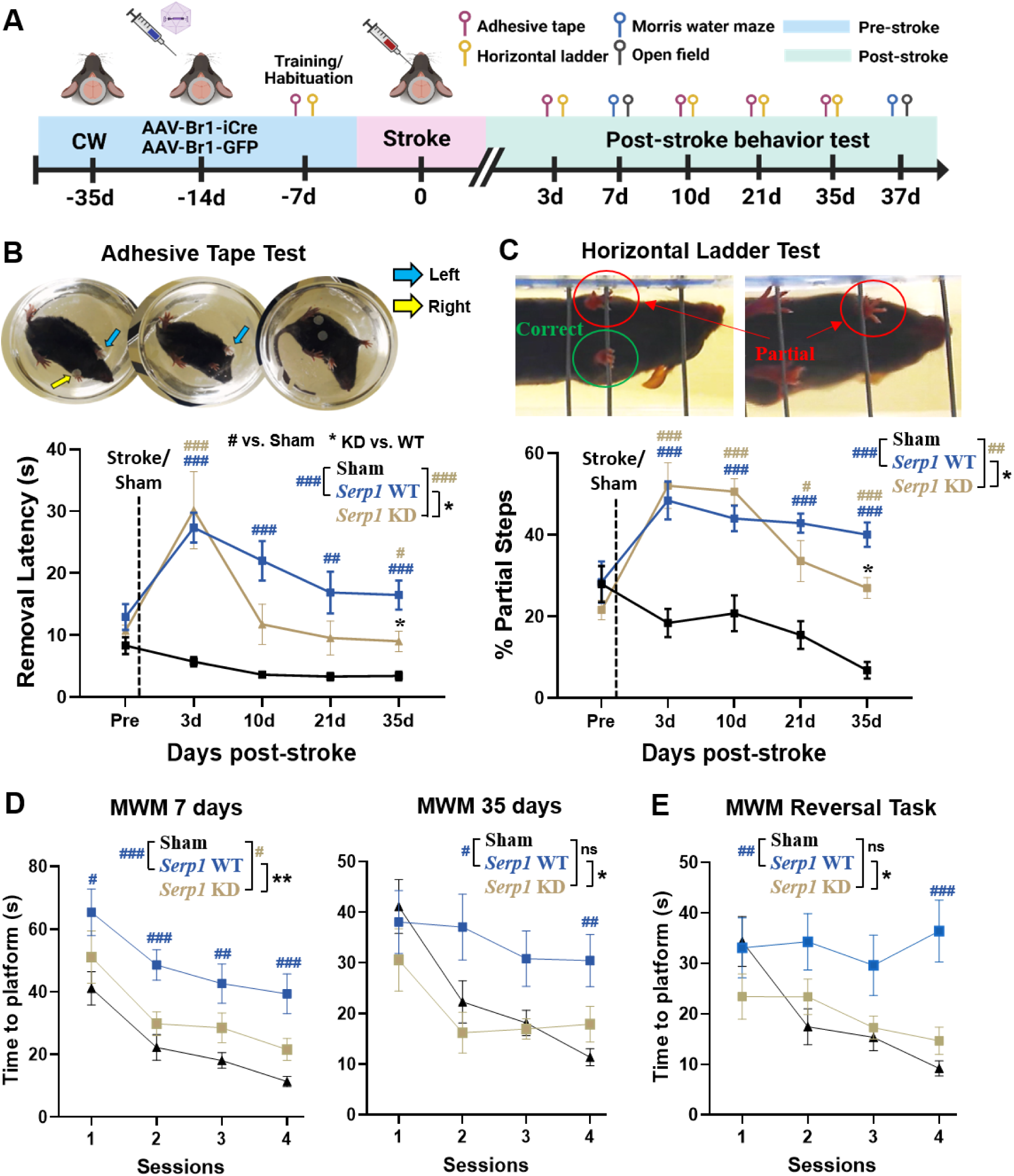
S*e*rpine1 KD mice show improved recovery of sensori-motor and cognitive function. **A.** Experimental timeline for behavioral tests. **B.** Representative example of a stroke affected mouse removing adhesive tape from left and right paws. Graph shows latency of tape removal (s) in the three groups; sham stroke (black, *n* = 15 mice), *Serp1* WT (blue, *n* = 13 mice) and *Serp1* KD (golden, *n* = 16 mice). **C.** Representative example of a stroke affected mouse crossing the horizontal ladder test, circles denote correct and partial paw placements. Graph shows average number of partial placement steps as a percent (%) total steps in the three experimental groups. **D.** Escape latency for learning the hidden platform location in Morris water maze (MWM) at 7d (left) and 35d (right) post-stroke. **E.** Average escape latency for learning new platform location in MWM (‘reversal task’) in the three groups. Data in **B-E** analyzed with two-way ANOVA and Tukey’s multiple comparison tests. Comparisons between *Serp1* KD vs WT: ** *p* < 0.01, * *p* < 0.05. Comparisons between each genotype and Pre-stroke or Sham stroke value: ### *p* < 0.001, ## *p* < 0.01, #p<0.05. Data expressed as the mean ± SEM.

## Discussion

In the present study we examined the role of endothelial *Serp1*/PAI-1 signalling, a critical component of the fibrinolysis cascade, in regulating cerebral microcirculation and recovery of brain function after ischemic stroke. Consistent with previous work, we show using immunohistochemistry and gene expression analysis that *Serp1*/PAI-1 is highly upregulated in peri-infarct regions after stroke ^11,12,14^. Using a viral strategy to knock down *Serp1* in endothelial cells, we unexpectedly found a significant decrease in capillary blood flow velocity and flux in peri-infarct cortex after stroke. Reduced blood flow was associated with attenuated BBB disruption and expression of inflammatory genes and cytokines at 3 days post-stroke. Over the longer term, endothelial *Serp1* knockdown led to improved recovery of sensory evoked cortical responses and behavioural measures of sensori-motor and cognitive function. Together, these findings show that endothelial specific manipulations of *Serp1*/PAI-1 signalling can potently modulate cerebral blood flow and improve functional recovery from stroke.

Optimal recovery from stroke is dependent on proper macro- and microvascular blood flow^4,9,45^. In the first few hours after an ischemic event, this undoubtedly means restoring adequate blood flow to allow vulnerable cells to survive ^6,46,47^. Case in point, alleviating the obstruction of large arteries with tPA or mechanical thrombectomy is standard clinical practice for improving stroke outcome. At the microvascular level, recent data show that preventing the stalling of peri-infarct capillaries in the first few hours after stroke, even after recanalizing large vessels with tPA, enhances return of cerebral blood flow in capillaries, reduces ischemic infarct volume, and behavioural metrics of neurological function ^48,49^. Over the longer term (days and weeks), what constitutes “optimal” blood flow in stroke affected regions, particular at the capillary level, is unclear. Conventional logic might suggest that increased capillary flow in peri-infarct regions would be better for stroke recovery. Indeed this is what motivated the present study and our knockdown approach since *Serp1*/PAI-1 signalling inhibits the breakdown of fibrin clots (see **Fig. 1A**), and therefore should increase post-stroke capillary blood flow. However as we found out, enhanced functional recovery in *Serp1* knockdown mice was associated with a reduction in capillary blood flow. Whether a reduction in peri-infarct capillary flow in *Serp1* KD mice at 3 days could be advantageous relative to the hyperemia found in WT mice is unknown, although modelling studies suggest that chronic hyperemia could impair capillary flow heterogeneity, which reduces oxygenation and maximum oxygen extraction factor in microvessels ^50,51^. It is important to note that enhanced recovery in *Serp1* knockdown mice could not be explained by an acute neuroprotective effect (ie. salvaging more penumbra) since infarct volumes were similar between experimental groups. This finding differs slightly from previous studies showing that systemic inhibition of PAI-1 or congenital knockout of *Serp1* improves stroke outcome by reducing the volume of infarction ^31,52,53^. However, neither previous study explored cell specific knockdown/inhibition approaches or examined functional recovery beyond 24 hours. Thus our study adds important new datum regarding endothelial specific manipulations of *Serp1*/PAI-1 signalling and its long-term effects on stroke recovery.

If *Serp1* knockdown does not provide any early neuroprotection, then how might it lead to better recovery? The fact that infarct volumes do not explain better recovery in *Serp1* knockdown mice is not surprising since infarct volume, especially when smaller in size, is considered a weak predictor of long term functional recovery^54,55^. Other factors, such as the BBB may be critical since it is disrupted for multiple days after a stroke which can directly disrupt neuronal structure and function in surviving peri-infarct regions and worsen stroke outcome^56,57^. Importantly, attenuating BBB disruption after stroke or in neurological diseases can rescue sensori-motor and cognitive deficits^39,58,59^. Previously, we showed that increased BBB permeability (at 3 days post-stroke) and poor functional outcome in diabetic mice, was accompanied by abnormally increased peri-infarct capillary blood flow relative to non-diabetic control mice^39^. We should also note that the non-diabetic control mice in the previous study still showed a transient increase in capillary flow at 3 days^9^, mirroring *Serp1* WT mice in the present work (see **Fig. 2E**). The idea that elevated blood flow in peri-infarct capillaries leads to increased BBB permeability (or vice versa) is plausible since these capillaries are in a vulnerable state with heightened remodelling (angiogenesis and pruning), and increased expression of ion channels and growth factors that modulate blood flow and BBB integrity^60–65^. Indeed we found that peri-infarct BBB permeability was lower in *Serp1* knockdown mice at 3 days recovery relative to WT mice. Lowered permeability was also associated with reduced expression of putative pro-inflammatory gene expression in peri-infarct regions. We did not find any group differences in BBB permeability at 10 and 21 days recovery although our sham stroke data suggest that the stroke induced disruption of the BBB has largely resolved by 10 days (Note in **Fig. 4B** that shams and stroke affected mice are similar at day 10 and 21). Our findings are reminiscent of work showing that returning blood flow too quickly in the hyper-acute stage of stroke with too few cerebral collaterals, leads to hemorrhages and worse stroke outcome in mice and humans^7,8^. Thus, our data suggest that a reduction in peri-infarct capillary blood flow could be beneficial for stroke recovery by attenuating BBB disruption.

A reduction in BBB permeability in *Serp1* KD mice may also promote recovery by lessening neuro-inflammation and potentially damaging processes associated with it. To explore this, we examined inflammatory gene expression in peri-infarct cortex at 3 days. GO pathway analysis revealed lowered expression of inflammation related pathways in *Serp1* KD mice relative to WT controls. For example, inflammatory genes that are prevalent after injury^15,66^ such as *Spp1*, *Slamf9*, *Lcn2*, *Top2a*, *Lilrb4a*, and *Tgm1* were downregulated in *Serp1* KD. Similarly, there was decreased expression of genes involved in chemokines (*Ccl2, 3, 4 5*) and the complement system (*C3/5ar1, C1qa/b/c, C1qb*) in the *Serp1* KD group ^13,67^. Conversely some anti-inflammatory and neuroprotective genes were increased in *Serp1* KD mice. These include *Uty*, a gene involved in protection against hypertension and reducing pro-inflammatory cytokines as well as *Pink1*, which protects against oxidative stress-induced apoptosis^68,69^. Blood serum analysis also revealed lowered expression of IL-6 in in *Serp1* KD mice. IL-6 is known to stimulate leukocyte activation/proliferation, and blood serum levels are predictive of worse stroke outcome and recurrent vascular diseases in human and animal models^70–73^. However, we are cautious of this simplistic explanation for improved recovery in *Serp1* KD mice and think future studies will be needed to unravel the complex interplay between PAI-1, inflammation and brain repair after stroke^74^.

While *Serp1*/PAI-1 signalling is best known for its role in inhibiting thrombolysis, there are reports suggesting that it can modulate vessel tone either directly or indirectly through tPA^23^. Our study is the first to show that endothelial knockdown of *Serp1* leads to constriction of peri-infarct capillaries. We found that capillaries branching off penetrating arterioles showed greatest constriction. Since these capillaries are enwrapped by either smooth muscle cells or pericytes^36,37,75^, it is possible that endothelial production of PAI-1 after stroke normally acts to relax vessel tone (ie. dilate vessels). This may explain why there was a transient increase in vessel diameter in WT mice 3 days after stroke, which replicates our previous data^9^. Of note, *Serp1* KD mice did show enhanced sensory evoked IOS cortical signals (**Fig. 6**), agreeing with past work showing improved neurovascular coupling (based on laser Doppler or speckle) following systemic inhibition of PAI-1 with genetic or pharmacological approaches^76,77^. Since IOS signals reflect fractional changes in de-oxyhemoglobin based absorption of red light, larger IOS responses in *Serp1* KD mice could reflect a greater dynamic range of blood flow response because narrower vessels (at baseline) would allow for greater increases in blood flow when stimulated. While the signalling mechanisms that couple PAI-1 with changes in vessel tone remain unclear, PAI-1 is highly enriched in myo-endothelial junctions^78^ and can interact with eNOS signalling^79,80^. Further PAI-1 can dose dependently stimulate arterial contractility^23^. In conclusion, our *in vivo* imaging data suggest that PAI-1 influences vessel tone at the level of capillaries, although more work will be needed to resolve the precise signalling mechanisms.

While our study provides novel pre-clinical evidence implicating endothelial *Serp1*/PAI-1 signalling as a determinant of stroke recovery, there are limitations to our study. First, we did not perform our imaging experiments in the hyper-acute stage of stroke (ie. <24 hours). This decision was guided by the fact that infarct volumes were similar between groups and that imaging peri-infarct tissues soon after stroke is technically challenging given substantial edema and tissue distortion^81^. Second, given the technical challenges that are inherent with imaging the same blood capillaries over many weeks after stroke, our study was not sufficiently powered to detect sex differences despite the fact that both male and female mice were sampled. Resolving potential sex differences is a critical issue but would require many more samples and thus better served in a future study. And finally, the translational significance of our study could be questioned since our cell specific knockdown was initiated before, not after stroke. However we are encouraged by new drug delivery systems that target endothelial cells^82^, thereby raising the prospect of a *Serp1*/PAI-1 inhibition strategy that could be initiated after stroke. These cell specific manipulations are an important consideration for future studies since it could avoid unwanted side effects such as hemorrhages that are usually associated with systemic manipulations of PAI-1. In this light, our study raises the possibility that endothelial knockdown or inhibition of *Serp1*/PAI-1 can improve stroke recovery without inducing vascular complications.

## Materials and Methods

### Animals

Adult (2–4 month old) female and male *Serpine1*^fl/fl^ mice on C57BLJ background (Jackson #032522) were used in all experiments. Although male and female mice were included in each experiment, we did not have sufficient sampling of each sex and statistic power to draw conclusions about possible sex differences. *Serpine1* floxed mice were bred to homozygosity (*Serpine1*^fl/fl^) and offspring were genotyped using the following primers: Wild-type; 5’ – CAC-CAT-GAA-AGG-GTA-CCA-CTG-TCA-G – 3’, Floxed; 5’ GCG-CCG-GAA-CCG-AAG-T 3’. Mice were housed under 12-hour light/dark cycle and given *ad litbitum* access to water and laboratory diet (Picolab Rodent diet 20, #50553). Ventilating housing racks were controlled for temperature (21 - 23°C) and humidity (40 – 50%) sensitive environment. All experiments were conducted according to the guidelines set by the Canadian Council of Animal Care and approved by the University of Victoria Animal Care Committee.

### Craniotomies

Mice were anaesthetized with isoflurane (2% induction and 1.2% maintenance) in medical grade air at a flow rate of 0.7 L/min and head-fixed to a custom-built surgical stage. Animal’s body temperature was maintained at 37°C throughout the procedure. Lidocaine was injected subcutaneously under the scalp at the surgical site, and 0.03mL bolus of 2% dexamethasone was injected intramuscularly to reduce inflammation. A midline incision was performed followed by scalp retraction. After teasing off soft tissues on the skull, a custom metal ring (outer diameter 11.3 mm, inner diameter 7.0 mm, height 1.5mm) was affixed to the skull with Metabond adhesive over the right somatosensory cortex (∼2mm right of bregma). Using a high-speed dental drill, two circular areas were thinned overlying the somatosensory cortex: the outer ring 6mm in diameter, and the inner ring 5mm in diameter. Cold (<4°C) HEPES-buffered artificial cerebrospinal fluid (ACSF) was applied periodically throughout to cool the exposed brain surface and reduce inflammation. The inner thinned area of the skull was removed with forceps leaving the dura intact and cold ACSF was left on the brain surface for 15min to reduce possible swelling. A custom glued cover slip (6mm outer, 5mm inner) was placed over the intact dura and secured to the skull with cyanoacrylate glue and dental cement. Mice were allowed recover on a heating pad before being returned to their cage.

### AAV based knockdown of *Serp1* in brain endothelial cells

We used the AAV-BR1 serotype for expressing Cre recombinase in endothelial cells of *Serp1*^fl/fl^ mice^34^. *Serp1*^fl/fl^ mice were anesthetized with ∼1.5% isofluorane in medical air and intravenously injected with 20µL of AAV-BR1-iCre (5.0×10^12^ GC/mL). Wild-type controls consisted of *Serp1*^fl/fl^ mice injected with equivalent doses of AAV-BR1-eGFP.

### Photothrombic stroke

We performed targeted ischemic stroke to the right forelimb (FL) somatosensory cortex (mapped from IOS imaging) using the photothrombotic method^33,83^. Mice with cranial windows previously installed were lightly anesthetized with ∼1.3% isofluorane in medical air and fitted to a custom imaging stage under an upright Olympus BX51WI microscope. Animal body temperature was maintained at 37°C throughout the procedure. Mice were administered 1% Rose Bengal solution (100mg/kg in HEPES-buffered ACSF) via intraperitoneal (i.p.) injection. The FL cortex was illuminated with a green LED (∼10mW, wavelength centered at 530nm) directed through a 10x objective lens for 15-20 min, or until the targeted surface vessel stopped flowing. Sham stroke group received only an i.p. injection of 1% Rose Bengal but no illumination.

### *In vivo* two-photon imaging of capillary blood flow and BBB permeability

Mice were lightly anesthetized with ∼1% isofluorane in medical air and fitted to a custom imaging stage. Mice were intravenously injected with 2% FITC dextran for labelling blood plasma (Sigma, 46945, molecular weight 70kDa, in 0.9% saline). High resolution images of cerebral vasculature were acquired using an Olympus FV1000MPE laser scanning microscope fed by a mode-locked Ti:Sapphire laser source with an average delivered laser power of 15 to 50mW at 800nm wavelength. Emitted light was split by a dichroic filter (552nm) before it passed through either a 495-540nm or 558-706nm bandpass filter. Image stacks were acquired to a depth of ∼250µm below the pial surface with a 20x Olympus XLUPlanFl water-immersion objective (NA=0.95) using Olympus Fluoview FV10-ASW software. Stacks were collected at zoom of 1.3, z-step size of 1.25μm, covering an area of 489.5×489.5µm (0.48μm/pixel). Cortical areas close to (range from 113 to 352µm) and further away (range from 491 to 1643µm) from the infarct border were imaged and relocated at respective days using local vessel landmarks and reference coordinates.

To measure RBC velocity, we collected line scans through the middle of several different capillary segments and repeated this 3 times with 30s between each scan to obtain the average RBC velocity for a capillary. Since moving RBCs are unlabeled, the resulting line scan has angled streaks, where the velocity is proportional to the slope of the RBC streaks. To automate our analysis of line scans, we employed particle image velocimetry (LS-PIV) MATLAB code^84^.

LS-PIV calculates the RBC velocity by determining the RBC displacement between pairs of the same line-scan using spatial cross-correlation analysis. This enabled us to measure a wide range of RBC velocities at varying conditions. We then convert the detected pixel shift (also the Δ distance in manual processing) to a velocity expressed in mm/s for each individual line scan. Vessel width was measured using the VasoMetrics macro on ImageJ/Fiji^85^. In general, we collected fluorescence intensity profiles at 5µm intervals through a capillary across each branch order starting from the 1^st^ branch to the 6^th^. Diameter was then estimated based on the average distance between the two half-maximum intensity points on the Gaussian-like intensity profile.

For assessing BBB permeability, mice were prepared for imaging as described above and intravenously injected with 75µL of 2.5% lower molecular weight FITC dextran (4kDa, Sigma, 46944, in 0.9% saline). Two-photon imaging (wavelength set to 800nm) started within 5 min after FITC-dextran injection. Image stacks consisted of 10 images (Kalman average = 2), collected at 10µm z-steps from a depth of 0-100µm from cortical surface. These stacks were taken under identical laser power/imaging conditions, every 2 minutes for up to 30min. FITC-dextran extravasation was measured based on fluorescence intensity on a 12 bit scale, by placing an ROI (15×15µm) in the extravascular space (minimum ∼5-8µm from the nearest vessel). Changes in extravascular fluorescence intensity at each time point were normalized to the first time point (0 min).

### IOS imaging and pulse oximetry

Intrinsic optical signal imaging (IOS) was used to non-invasively and indirectly examine neural activity in primary somatosensory cortex. Mice with cranial windows previously installed were lightly anesthetized with ∼1% isofluorane in medical air and fitted to a custom imaging stage. A pencil lead (0.7mm) connected to a piezoelectric bending actuator (Piezo Systems, Q220-A4-203YB; 300μm deflection) was lightly glued to the FL and HL separately. The cortical surface was illuminated with a red LED (625nm) coupled to a collimation lens. Sensory-evoked changes in reflected light were collected for FL and HL (separately) using the MiCAM-2 CCD camera (SciMedia) through a 2X objective lens. Each imaging trial was collected over 3s at 100Hz with a 10ms exposure. Vibrotactile stimulation (100Hz) for each limb was initiated after 1s in each 3s trial and lasted for 1s. Two sets of 12 sensory stimulation trials were recorded for each limb. Using FIJI software, the 24 trials were averaged and then mean filtered (radius 5 pixels). Each image in the averaged and smoothed image stack was then subtracted and divided from a mean intensity projection of the pre-stimulus period (frames from 0-1s).

Delineation of FL and HL regions was based on thresholding an average intensity image projection (projected frames from 0.5-2s after stimulation) at half-maximal intensity. IOS signals were then measured by placing an ROI (0.5mm diameter) in the center of the IOS response. Peak amplitudes for FL and HL responses were then measured using Clampfit software.

Physiological parameters such as O2 saturation, heart and breath rate were measured during IOS imaging sessions before stroke and 3 days post-stroke. A tail cuff was placed onto the mouse and measurements were recorded at 10Hz using the MouseOx small animal vital signs monitor (Starr Life Sciences) for a total of 15-20min at each time point.

### Immunofluorescence and confocal imaging

Mice were administered an overdose of sodium pentobarbital and transcradially perfused using cold phosphate-buffered saline (PBS). Brains were extracted and then either frozen in dry ice or fixed overnight in 4% PFA followed by 30% sucrose solution in 0.1M PBS with 0.2% sodium azide. Fresh frozen brains were sectioned on a cryostat at 40µm and fixed in ice cold methanol. PFA fixed brains were sectioned at 40µm on a freezing microtome (American Optical Corp.) and stored in 0.1M PBS with 0.2% sodium azide solution. For antigen retrieval, sections were incubated in sodium citrate buffer (10mM, pH = 6.0) for 30min at 75°C. Following a wash in 0.1M PBS for 5 mins, sections were incubated with PAI-1 primary antibody (monoclonal mouse; 1:1000, Abcam) overnight at 4°C. Sections were washed with PBS followed by incubation with secondary antibody (Cy5 or Alexa Fluor 568 conjugated anti-mouse; 1:500, Invitrogen) for 4 hrs at room temperature. Sections were mounted, coverslipped and imaged using an Olympus confocal microscope at 10x (NA = 0.40) and 20x objective (NA = 0.75). Fluorophores Cy5 and/or 568 were excited at 615 and 568nm lasers, respectively. Image stacks were collected in 2 (for 10x obj. lens) or 1 (for 20x obj. lens) µm Z-steps at a resolution of 1.24 and 0.621µm/pixel, respectively. For vessel analysis, stacks were maximally z-projected and automatically thresholded using built in ImageJ/Fiji threshold function Triangle^86^, which best identified vessel signal. A median filter (radius 1 pixel) was applied to eliminate speckling artifact and the remaining area of vascular signal was measured within the peri-infarct region, and then normalized to sham stroke controls.

### Behavioural testing

Adhesive tape removal and horizontal ladder tests were included for their high sensitivity in detecting damage-related changes in sensory-motor function of the forepaw^87–89^. To assess spatial learning and memory we performed Morris Water Maze^90^ as well as a reversal learning test for cognitive flexibility^91^. And to assess locomotive ability, we performed open-field test ^92^. Behavioural tests were administered 2 weeks before stroke and at each two-photon imaging time-point afterwards; behavioral tests were conducted in the morning, and imaging in the evening. All of the following tests are described from previous work in the lab ^39,44,88,93^

#### Tape Removal Test

A circular piece of tape (5 mm diameter) was adhered to the palmar surface of each forepaw. Mice were then placed in a clear glass cylinder and filmed from below for up to 120s until the tape pieces were removed from both forepaws. The latency to remove the tape from each forepaw was recorded in seconds and both forepaws were averaged as tape removal latency (s). This was repeated three times per testing session.

#### Horizontal Ladder Test

Mice were videotaped walking across an elevated 70-cm-long horizontal ladder that had 1 mm diameter rungs randomly spaced 1 or 2 cm apart. Grasping of the rungs by the forepaws were scored on a frame-by-frame basis based on the centrality and completeness of the forepaw placement. The scores were assigned by a blind observer as follows: (i) “correct” (forepaw placement centered), (ii) “partial” (forepaw placement partially centered), (iii) “slip/miss” (forepaw placement not centered).

#### Open field test

Mice were placed into the center of a 100 cm diameter circular arena, and the total distance travelled (cm) was automatically recorded and measured using Ethovision software (Noldus Information Technology, Version 11.5.1020, 2015). One trial of 7min was conducted for each mouse 2 weeks pre-stroke, and 7-and 35-days post-stroke.

#### Morris Water Maze

In a large circular water-filled pool (100 cm diameter), a hidden platform was placed approximately 1 cm below the surface in one of four quadrants. Non-toxic white paint was added to the water to hide the platform, and water temperature was maintained at room temp. Mice were trained to locate the hidden platform using unique shaped-visual cues (maintained throughout experiment) placed around the maze. Mice were then placed in one quadrant facing a specified starting point: northeast (NE), southeast (SE), northwest (NW), or northeast (NE). The trial ended when the platform was located and mice stopped on the platform for three seconds, or until a maximum of two minutes had elapsed without the platform being located. The training sessions were performed over 2 days for a total of four training sessions. To score escape latency (seconds) and swim speeds (cm/s) to platform, the Ethovision software was used. For the reversal task, we switched the platform to a new location and performed the test in the same manner as the initial training and trial sessions.

### Nanostring and cytokine multiplex assays

In brief, mice were deeply anaesthetized and perfused with cold, sterile 0.1M PBS for 30 sec before tissue extraction. Peri-infarct regions in stroke mice or equivalent regions in sham controls (*n* = 4 per group) were isolated and flash-frozen in liquid nitrogen and stored at -80°C. A minimum of approximately 100ng of total RNA was used, and RNA quality and quantity (A260/A280) were estimated using Nanodrop 200 spectrophotometer (ThermoFisher, ND-2000). Samples were processed using the NanoString nCounter mouse neuroinflammation panel (770 genes + 6 internal reference controls; Catalogue IDL XT_Mm_Neuroinflam_CSO – 115000237, Nanostring Technologies, Seattle, WA, USA). Hybridization was performed using nCounter Master Kit (Catalogue # NAA-AKIT-012, Bruker). The number of occurrences of the hybridized cDNA to the unique bar-carded probes was counted by Nanostring nCounter instrument and raw gene counts were obtained for further analysis. Nanostring nSolver version 4.0 was used to extract raw gene counts for all tissue samples. All experimental samples were organized based on their condition; Sham stroke, *Serp1* WT stroke, and *Serp1* KD stroke. Background threshold was calculated as 12 counts, using the ‘mean negative probe counts + 2 [standard deviation of negative probe counts]’ formula, and gene counts < 12 were excluded from the analysis. Gene counts were then normalized to positive control and housekeeping probes, and resulting normalized and annotated data was used for subsequent analysis.

For examining cytokine levels in blood serum, mice were deeply anaesthetized and blood was collected and allowed to clot at room temperature for 10min and then centrifuged at 1500 rpm for 10 min at 4°C. The supernatant was collected and diluted 2-fold using standard PBS pH ∼7.5 before flash-frozen in liquid nitrogen and stored at -80°C. Serum levels (in duplicates) of the cytokines were determined using Luminex 200 platform and mouse cytokine/chemokine 32-plex discovery assay (MD32) by Eve Technologies (Calgary, AB, Canada). Raw values for each group were normalized to sham stroke controls.

### Statistics

Statistical analysis of the data was conducted using R/R-studio (Version 4.4.1) and GraphPad Prism (version 10.4.2). One-way analysis of variance (ANOVA) was used to compare baseline measurements of vascular parameters (blood flow and vessel width) across experimental groups. Planned two sample unpaired t-tests were used for comparing groups of 3 or less (**Fig. 1E-G**, **Fig. 2A**). One-way ANOVA followed by Sidak comparisons was used for analyzing PAI-1 staining over time. Two-way ANOVA was used to analyze effects of stroke, genotype (*Serp1* knockdown), or interaction effects on RBC velocity/flux, vessel width, BBB leakage, IOS responses, cytokine panel and behaviour. Significant main effects or interactions were followed up by *post-hoc* multiple comparisons (Tukey, or Mann-Whitney on the data set). A moderated t-test with multiple test corrections using limma was used to determine significance in gene expression. Transcriptomic plots including volcano plots, heatmaps, horizontal bar graphs, and line plots were made using R/R-studio. RBC velocities from line scans were acquired using MathLab R2023a. All data are presented as mean ± standard error of mean (SEM).

## Acknowledgments

We thank A. Hentze, T. Yang, and ACU staff for managing the mouse colony. We also thank Dr. P. Reeson, and Dr. S. Sharma for thoughtful discussions regarding the work.

## References

1. Joy, M. T. & Carmichael, S. T. Encouraging an excitable brain state: mechanisms of brain repair in stroke. Nat. Rev. Neurosci. 22, 38–53 (2021).

2. Clark, T. A. et al. Artery targeted photothrombosis widens the vascular penumbra, instigates peri-infarct neovascularization and models forelimb impairments. Sci. Rep. 9, 2323 (2019).

3. Schrandt, C. J., Kazmi, S. S., Jones, T. A. & Dunn, A. K. Chronic Monitoring of Vascular Progression after Ischemic Stroke Using Multiexposure Speckle Imaging and Two-Photon Fluorescence Microscopy. J. Cereb. Blood Flow Metab. 35, 933–942 (2015).

4. Zhang, S., Boyd, J., Delaney, K. & Murphy, T. H. Rapid Reversible Changes in Dendritic Spine Structure In Vivo Gated by the Degree of Ischemia. J. Neurosci. 25, 5333–5338 (2005).

5. Schaffer, C. B. et al. Two-Photon Imaging of Cortical Surface Microvessels Reveals a Robust Redistribution in Blood Flow after Vascular Occlusion. PLOS Biol. 4, e22 (2006).

6. Shih, A. Y. et al. Active Dilation of Penetrating Arterioles Restores Red Blood Cell Flux to Penumbral Neocortex after Focal Stroke. J. Cereb. Blood Flow Metab. 29, 738–751 (2009).

7. Binder, N. F. et al. Leptomeningeal collaterals regulate reperfusion in ischemic stroke and rescue the brain from futile recanalization. Neuron 112, 1456–1472.e6 (2024).

8. Yang, W. et al. Global hyperperfusion after successful endovascular thrombectomy is linked to worse outcome in acute ischemic stroke. Sci. Rep. 14, 10024 (2024).

9. Tennant, K. A. & Brown, C. E. Diabetes Augments In Vivo Microvascular Blood Flow Dynamics after Stroke. J. Neurosci. 33, 19194–19204 (2013).

10. Yanev, P. et al. Plasminogen activator inhibitor-1 mediates cerebral ischemia-induced astrocytic reactivity. J. Cereb. Blood Flow Metab. 45, 102–114 (2025).

11. Garcia-Bonilla, L. et al. Analysis of brain and blood single-cell transcriptomics in acute and subacute phases after experimental stroke. Nat. Immunol. 25, 357–370 (2024).

12. Munji, R. N. et al. Profiling the mouse brain endothelial transcriptome in health and disease models reveals a core blood-brain barrier dysfunction module. Nat. Neurosci. 22, 1892– 1902 (2019).

13. Hammond, T. R. et al. Single-Cell RNA Sequencing of Microglia throughout the Mouse Lifespan and in the Injured Brain Reveals Complex Cell-State Changes. Immunity 50, 253–271.e6 (2019).

14. Callegari, K., et al. Molecular profiling of the stroke-induced alterations in the cerebral microvasculature reveals promising therapeutic candidates. Proc. Natl. Acad. Sci. 120, e2205786120 (2023).

15. Weber, R. Z. et al. A molecular brain atlas reveals cellular shifts during the repair phase of stroke. J. Neuroinflammation 22, 112 (2025).

16. Akhter, M. S., Biswas, A., Abdullah, S. M., Behari, M. & Saxena, R. The Role of PAI-1 4G/5G Promoter Polymorphism and Its Levels in the Development of Ischemic Stroke in Young Indian Population. Clin. Appl. Thromb. 23, 1071–1076 (2017).

17. Cesari, M., Pahor, M. & Incalzi, R. A. PLASMINOGEN ACTIVATOR INHIBITOR-1 (PAI-1): A KEY FACTOR LINKING FIBRINOLYSIS AND AGE-RELATED SUBCLINICAL AND CLINICAL CONDITIONS. Cardiovasc. Ther. 28, e72–e91 (2010).

18. Chan, S.-L., Bishop, N., Li, Z. & Cipolla, M. J. Inhibition of PAI (Plasminogen Activator Inhibitor)-1 Improves Brain Collateral Perfusion and Injury After Acute Ischemic Stroke in Aged Hypertensive Rats. Stroke 49, 1969–1976 (2018).

19. Naya, M. et al. Elevated Plasma Plasminogen Activator Inhibitor Type-1 is an Independent Predictor of Coronary Microvascular Dysfunction in Hypertension. Circ. J. 71, 348– 353 (2007).

20. Tjärnlund-Wolf, A., Brogren, H., Lo, E. H. & Wang, X. Plasminogen Activator Inhibitor-1 and Thrombotic Cerebrovascular Diseases. Stroke J. Cereb. Circ. 43, 2833–2839 (2012).

21. Hamsten, A. et al. PLASMINOGEN ACTIVATOR INHIBITOR IN PLASMA: RISK FACTOR FOR RECURRENT MYOCARDIAL INFARCTION. The Lancet 330, 3–9 (1987).

22. Hamsten Anders, Wiman Björn, de Faire Ulf, & Blombäck Margareta. Increased Plasma Levels of a Rapid Inhibitor of Tissue Plasminogen Activator in Young Survivors of Myocardial Infarction. N. Engl. J. Med. 313, 1557–1563 (1985).

23. Nassar, T. et al. In vitro and in vivo effects of tPA and PAI-1 on blood vessel tone. Blood 103, 897–902 (2004).

24. Nassar, T. et al. tPA regulates pulmonary vascular activity through NMDA receptors. Am. J. Physiol. Lung Cell. Mol. Physiol. 301, L307–314 (2011).

25. Nagai, N., De Mol, M., Lijnen, H. R., Carmeliet, P. & Collen, D. Role of Plasminogen System Components in Focal Cerebral Ischemic Infarction. Circulation 99, 2440–2444 (1999).

26. Nagai, N., Suzuki, Y., Van hoef, B., Lijnen, H. R. & Collen, D. Effects of plasminogen activator inhibitor-1 on ischemic brain injury in permanent and thrombotic middle cerebral artery occlusion models in mice. J. Thromb. Haemost. 3, 1379–1384 (2005).

27. Eren, M., Painter, C. A., Atkinson, J. B., Declerck, P. J. & Vaughan, D. E. Age-Dependent Spontaneous Coronary Arterial Thrombosis in Transgenic Mice That Express a Stable Form of Human Plasminogen Activator Inhibitor-1. Circulation 106, 491–496 (2002).

28. Erickson, L. A. et al. Development of venous occlusions in mice transgenic for the plasminogen activator inhibitor-1 gene. Nature 346, 74–76 (1990).

29. Griemert, E.-V., Recarte Pelz, K., Engelhard, K., Schäfer, M. K. & Thal, S. C. PAI-1 but Not PAI-2 Gene Deficiency Attenuates Ischemic Brain Injury After Experimental Stroke. Transl. Stroke Res. 10, 372–380 (2019).

30. Sashindranath, M. et al. The tissue-type plasminogen activator–plasminogen activator inhibitor 1 complex promotes neurovascular injury in brain trauma: evidence from mice and humans. Brain 135, 3251–3264 (2012).

31. Palakurti, R. et al. Inducible miR-1224 silences cerebrovascular Serpine1 and restores blood flow to the stroke-affected site of the brain. Mol. Ther. Nucleic Acids 31, 276–292 (2023).

32. Denorme, F., et al. Inhibition of Thrombin-Activatable Fibrinolysis Inhibitor and Plasminogen Activator Inhibitor-1 Reduces Ischemic Brain Damage in Mice. Stroke 47, 2419–2422 (2016).

33. Watson, B. D., Dietrich, W. D., Busto, R., Wachtel, M. S. & Ginsberg, M. D. Induction of reproducible brain infarction by photochemically initiated thrombosis. Ann. Neurol. 17, 497–504 (1985).

34. Körbelin, J. et al. A brain microvasculature endothelial cell-specific viral vector with the potential to treat neurovascular and neurological diseases. EMBO Mol. Med. 8, 609–625 (2016).

35. Jin, Y. et al. Longitudinal, Multimodal Tracking Reveals Lasting Neurovascular Impact of Individual Microinfarcts. Adv. Sci. 12, 2417003 (2025).

36. Hall, C. N. et al. Capillary pericytes regulate cerebral blood flow in health and disease. Nature 508, 55–60 (2014).

37. Hartmann, D. A. et al. Brain capillary pericytes exert a substantial but slow influence on blood flow. Nat. Neurosci. 24, 633–645 (2021).

38. Blinder, P. et al. The cortical angiome: an interconnected vascular network with noncolumnar patterns of blood flow. Nat. Neurosci. 16, 889–897 (2013).

39. Reeson, P. et al. Delayed Inhibition of VEGF Signaling after Stroke Attenuates Blood– Brain Barrier Breakdown and Improves Functional Recovery in a Comorbidity-Dependent Manner. J. Neurosci. 35, 5128–5143 (2015).

40. Korte, N. et al. Inhibiting Ca2+ channels in Alzheimer’s disease model mice relaxes pericytes, improves cerebral blood flow and reduces immune cell stalling and hypoxia. Nat. Neurosci. 27, 2086–2100 (2024).

41. Kutuzov, N., Flyvbjerg, H. & Lauritzen, M. Contributions of the glycocalyx, endothelium, and extravascular compartment to the blood-brain barrier. Proc. Natl. Acad. Sci. U. S. A. 115, E9429–E9438 (2018).

42. Zeiger, W. A. et al. Barrel cortex plasticity after photothrombotic stroke involves potentiating responses of pre-existing circuits but not functional remapping to new circuits. Nat. Commun. 12, 3972 (2021).

43. Bandet, M. V. & Winship, I. R. Aberrant cortical activity, functional connectivity, and neural assembly architecture after photothrombotic stroke in mice. eLife 12, RP90080 (2024).

44. Motaharinia, M. et al. Longitudinal functional imaging of VIP interneurons reveals sup-population specific effects of stroke that are rescued with chemogenetic therapy. Nat. Commun. 12, 6112 (2021).

45. Williamson, M. R. et al. A Window of Vascular Plasticity Coupled to Behavioral Recovery after Stroke. J. Neurosci. 40, 7651–7667 (2020).

46. Dalkara, T. & Arsava, E. M. Can restoring incomplete microcirculatory reperfusion improve stroke outcome after thrombolysis? J. Cereb. Blood Flow Metab. 32, 2091–2099 (2012).

47. Murphy, T. H. & Corbett, D. Plasticity during stroke recovery: from synapse to behaviour. Nat. Rev. Neurosci. 10, 861–872 (2009).

48. Amki, M. E. et al. Neutrophils Obstructing Brain Capillaries Are a Major Cause of No-Reflow in Ischemic Stroke. Cell Rep. 33, (2020).

49. Erdener, Ş. E., et al. Dynamic capillary stalls in reperfused ischemic penumbra contribute to injury: A hyperacute role for neutrophils in persistent traffic jams. J. Cereb. Blood Flow Metab. 41, 236–252 (2021).

50. Jespersen, S. N. & Østergaard, L. The roles of cerebral blood flow, capillary transit time heterogeneity, and oxygen tension in brain oxygenation and metabolism. J. Cereb. Blood Flow Metab. 32, 264–277 (2012).

51. Østergaard, L. et al. Capillary transit time heterogeneity and flow-metabolism coupling after traumatic brain injury. J. Cereb. Blood Flow Metab. 34, 1585–1598 (2014).

52. Xu, Y. et al. Specific inhibition on PAI-1 reduces the dose of Alteplase for ischemic stroke treatment. Int. J. Biol. Macromol. 257, 128618 (2024).

53. Torrente, D. et al. Compartmentalized actions of the plasminogen activator inhibitors, PAI-1 and Nsp in ischemic stroke. Transl. Stroke Res. 13, 801–815 (2022).

54. Ospel, J. M. et al. Strength of Association between Infarct Volume and Clinical Outcome Depends on the Magnitude of Infarct Size: Results from the ESCAPE-NA1 Trial. AJNR Am. J. Neuroradiol. 42, 1375–1379 (2021).

55. Morais, A. et al. Biological and Procedural Predictors of Outcome in the Stroke Preclinical Assessment Network (SPAN) Trial. Circ. Res. 135, 575–592 (2024).

56. Venkat, P., Chopp, M. & Chen, J. Blood–Brain Barrier Disruption, Vascular Impairment, and Ischemia/Reperfusion Damage in Diabetic Stroke. J. Am. Heart Assoc. 6, e005819 (2017).

57. Geng, J. et al. Blood-brain barrier disruption induced cognitive impairment is associated with increase of inflammatory cytokine. Front. Aging Neurosci. 10, 1–11 (2018).

58. Montagne, A. et al. Blood-Brain Barrier Breakdown in the Aging Human Hippocampus. Neuron 85, 296–302 (2015).

59. Shi, S. M. et al. Glycocalyx dysregulation impairs blood–brain barrier in ageing and disease. Nature 639, 985–994 (2025).

60. Rust, R. Insights into the dual role of angiogenesis following stroke. J. Cereb. Blood Flow Metab. 40, 1167–1171 (2020).

61. Ohab, J. J., Fleming, S., Blesch, A. & Carmichael, S. T. A Neurovascular Niche for Neurogenesis after Stroke. J. Neurosci. 26, 13007–13016 (2006).

62. Lim, X. R. et al. Endothelial Piezo1 channel mediates mechano-feedback control of brain blood flow. Nat. Commun. 15, 8686 (2024).

63. Harraz, O. F., Klug, N. R., Senatore, A. J., Hill-Eubanks, D. C. & Nelson, M. T. Piezo1 Is a Mechanosensor Channel in Central Nervous System Capillaries. Circ. Res. 130, 1531–1546 (2022).

64. Longden, T. A., Hill-Eubanks, D. C. & Nelson, M. T. Ion channel networks in the control of cerebral blood flow. J. Cereb. Blood Flow Metab. 36, 492–512 (2016).

65. Kim, E., Yang, J., Park, K. W. & Cho, S. Inhibition of VEGF Signaling Reduces Diabetes-Exacerbated Brain Swelling, but Not Infarct Size, in Large Cerebral Infarction in Mice. Transl. Stroke Res. 9, 540–548 (2018).

66. Jin, C., et al. Leveraging single-cell RNA sequencing to unravel the impact of aging on stroke recovery mechanisms in mice. Proc. Natl. Acad. Sci. 120, e2300012120 (2023).

67. Hickman, S. E. et al. The Microglial Sensome Revealed by Direct RNA Sequencing. Nat. Neurosci. 16, 1896–1905 (2013).

68. Barodia, S. K., Creed, R. B. & Goldberg, M. S. Parkin and PINK1 functions in oxidative stress and neurodegeneration. Brain Res. Bull. 133, 51–59 (2017).

69. Cunningham, C. M. et al. Y-Chromosome Gene, Uty, Protects Against Pulmonary Hypertension by Reducing Proinflammatory Chemokines. Am. J. Respir. Crit. Care Med. 206, 186–196 (2022).

70. Tanaka, T., Narazaki, M. & Kishimoto, T. IL-6 in Inflammation, Immunity, and Disease. Cold Spring Harb. Perspect. Biol. 6, a016295 (2014).

71. Sharma, S. et al. A pathogenic role for IL-10 signalling in capillary stalling and cognitive impairment in type 1 diabetes. Nat. Metab. 6, 2082–2099 (2024).

72. Steen, E. H. et al. The Role of the Anti-Inflammatory Cytokine Interleukin-10 in Tissue Fibrosis. Adv. Wound Care 9, 184–198 (2020).

73. Barabási, B. et al. Role of interleukin-6 and interleukin-10 in morphological and functional changes of the blood-brain barrier in hypertriglyceridemia. Fluids Barriers CNS 20, 15 (2023).

74. Iadecola, C., Buckwalter, M. S. & Anrather, J. Immune responses to stroke: mechanisms, modulation, and therapeutic potential. J. Clin. Invest. 130, 2777–2788 (2020).

75. Hill, R. A. et al. Regional Blood Flow in the Normal and Ischemic Brain Is Controlled by Arteriolar Smooth Muscle Cell Contractility and Not by Capillary Pericytes. Neuron 87, 95–110 (2015).

76. Furon, J. et al. Blood tissue Plasminogen Activator (tPA) of liver origin contributes to neurovascular coupling involving brain endothelial N-Methyl-D-Aspartate (NMDA) receptors. Fluids Barriers CNS 20, 11 (2023).

77. Park, L. et al. tPA Deficiency Underlies Neurovascular Coupling Dysfunction by Amyloid-β. J. Neurosci. 40, 8160–8173 (2020).

78. Heberlein, K. R. et al. Plasminogen Activator Inhibitor-1 Regulates Myoendothelial Junction Formation. Circ. Res. 106, 1092–1102 (2010).

79. Goglia, L. et al. Endothelial regulation of eNOS, PAI-1 and t-PA by testosterone and dihydrotestosterone in vitro and in vivo. Mol. Hum. Reprod. 16, 761–769 (2010).

80. Garcia, V. et al. Unbiased proteomics identifies plasminogen activator inhibitor-1 as a negative regulator of endothelial nitric oxide synthase. Proc. Natl. Acad. Sci. 117, 9497–9507 (2020).

81. Kirov, S. A., Fomitcheva, I. V. & Sword, J. Rapid Neuronal Ultrastructure Disruption and Recovery during Spreading Depolarization-Induced Cytotoxic Edema. Cereb. Cortex N. Y. NY 30, 5517–5531 (2020).

82. Kucharz, K. et al. Post-capillary venules are the key locus for transcytosis-mediated brain delivery of therapeutic nanoparticles. Nat. Commun. 12, 4121 (2021).

83. Brown, C. E., Li, P., Boyd, J. D., Delaney, K. R. & Murphy, T. H. Extensive Turnover of Dendritic Spines and Vascular Remodeling in Cortical Tissues Recovering from Stroke. J. Neurosci. 27, 4101–4109 (2007).

84. Kim, T. N. et al. Line-Scanning Particle Image Velocimetry: An Optical Approach for Quantifying a Wide Range of Blood Flow Speeds in Live Animals. PLOS ONE 7, e38590 (2012).

85. McDowell, K. P., Berthiaume, A.-A., Tieu, T., Hartmann, D. A. & Shih, A. Y. VasoMetrics: unbiased spatiotemporal analysis of microvascular diameter in multi-photon imaging applications. Quant. Imaging Med. Surg. 11, 969–982 (2021).

86. Zack, G. W., Rogers, W. E. & Latt, S. A. Automatic measurement of sister chromatid exchange frequency. J. Histochem. Cytochem. 25, 741–753 (1977).

87. Shanina, E. V., Schallert, T., Witte, O. W. & Redecker, C. Behavioral recovery from unilateral photothrombotic infarcts of the forelimb sensorimotor cortex in rats: Role of the contralateral cortex. Neuroscience 139, 1495–1506 (2006).

88. Sweetnam, D. et al. Diabetes Impairs Cortical Plasticity and Functional Recovery Following Ischemic Stroke. J. Neurosci. 32, 5132–5143 (2012).

89. Tennant, K. A. & Jones, T. A. Sensorimotor Behavioral Effects of Endothelin-1 Induced Small Cortical Infarcts in C57BL/6 Mice. J. Neurosci. Methods 181, 18–26 (2009).

90. Morris, R. Developments of a water-maze procedure for studying spatial learning in the rat. J. Neurosci. Methods 11, 47–60 (1984).

91. Vorhees, C. V. & Williams, M. T. Morris water maze: procedures for assessing spatial and related forms of learning and memory. Nat. Protoc. 1, 848–858 (2006).

92. Seibenhener, M. L. & Wooten, M. C. Use of the Open Field Maze to Measure Locomotor and Anxiety-like Behavior in Mice. J. Vis. Exp. JoVE 52434 (2015) doi:10.3791/52434.

93. Brown, C. E., Aminoltejari, K., Erb, H., Winship, I. R. & Murphy, T. H. In Vivo Voltage-Sensitive Dye Imaging in Adult Mice Reveals That Somatosensory Maps Lost to Stroke Are Replaced over Weeks by New Structural and Functional Circuits with Prolonged Modes of Activation within Both the Peri-Infarct Zone and Distant Sites. J. Neurosci. 29, 1719–1734 (2009).

