## Supplemental data for "Knockdown of endothelial *Serpine1* improves stroke recovery by attenuating peri-infarct blood flow and blood brain barrier disruption"

Kamal Narayana *et al.*

### Supplementary Figures

#### SUPPLEMENTARY FIGURE 1

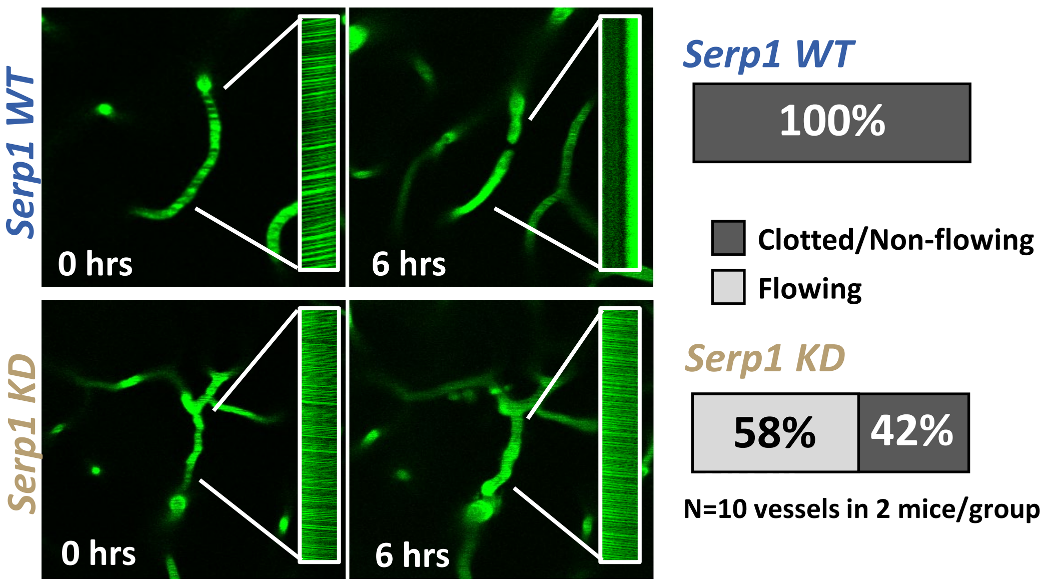


**Supp Fig. 1.** In vivo two-photon images and line scans of a vessel pre- and post-bleed (ie. 6hrs post-bleed) in *Serp1* WT and KD group. In *Serp1* WT group, all (100%) of the vessels were stalled at 6 hours, whereas, only 42% were stalled and 58% were flowing in Serp1 KD group.

#### SUPPLEMENTARY FIGURE 2

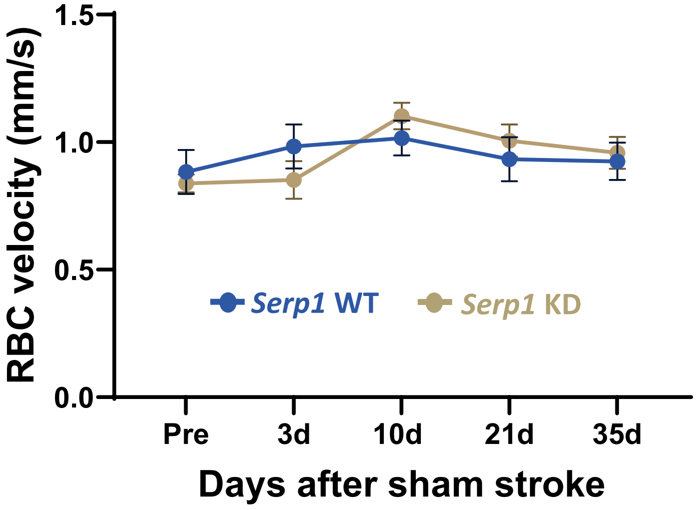


**Supp Fig. 2.** The sham stroke control group consisted of both *Serp1* WT and KD mice (n=3 from each genotype). RBC velocities did not differ between these groups (2-way ANOVA, Main effect of Genotype: $F_{(1, 560)}$ = 0.004, *=0.94*; Main effect of Time: $F_{(4, 560)}$ = 2.11 *p* = 0.07; Genotype x Time Interaction: $F_{(4, 560)}$ = 0.84, *p*=0.50). Data expressed as the mean ± SEM.

### SUPPLEMENTARY FIGURE 3

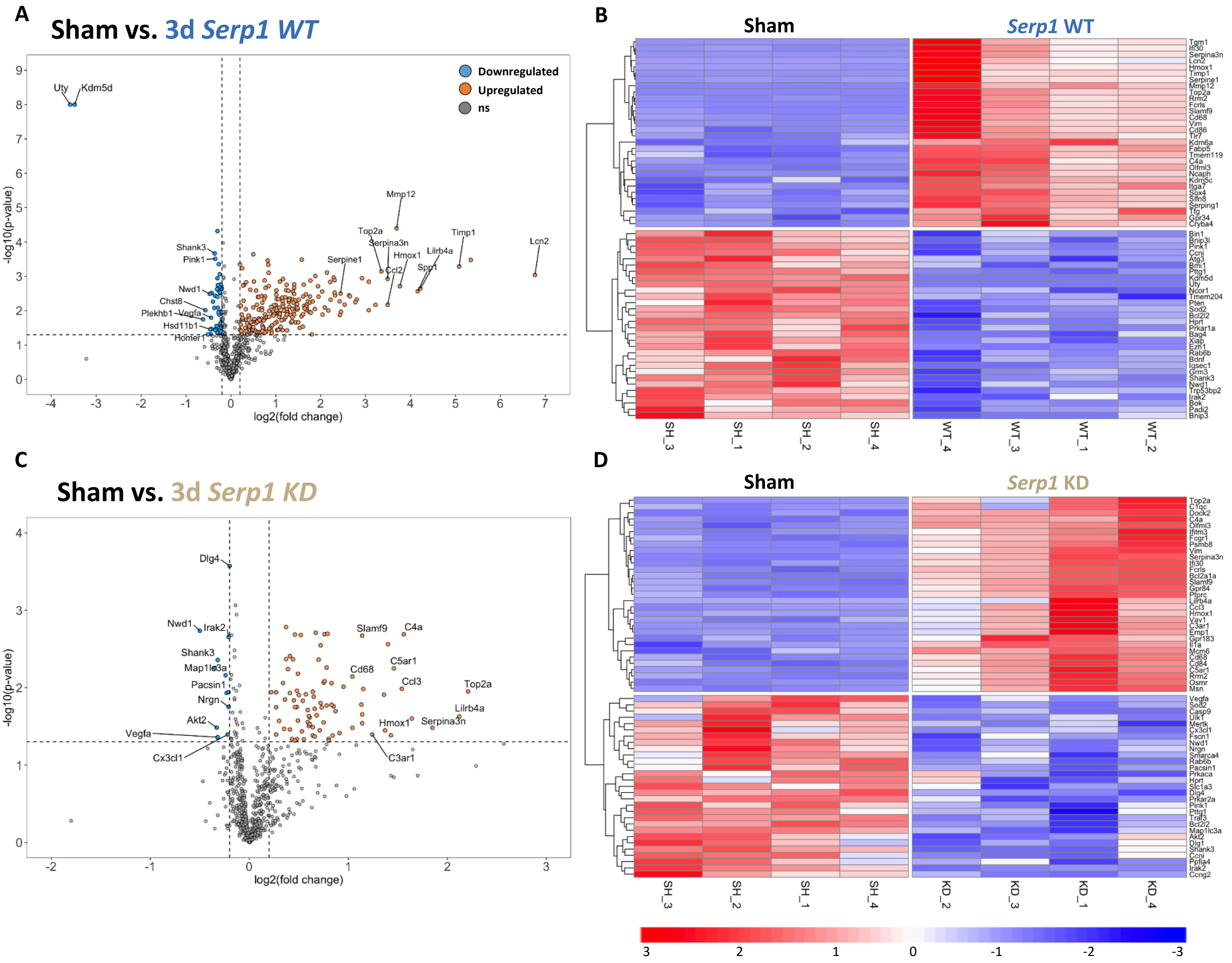


**Supp Fig. 3.** Transcriptomic changes in *Serpine1* WT and KD reveal intricate play of complex genes post-stroke. Volcano plot depicting differentially expressed genes (DEG) in **A.** *Serp1* WT vs. sham, and **C.** *Serp1* KD vs. sham as a function of $\log_2$ fold change (fc) and statistical significance of $p < 0.05$, denoted in y-axis as $-\log_{10}$($p$-value). Downregulated in blue, upregulated in orange, and non-significant in grey (and shaded). Top 10 up- and down-regulated and significant genes labelled. Heatmap (corresponding to volcano plot) of the top 30 regulated DEG in **B.** *Serp1* WT vs. sham, and **D.** *Serp1* KD vs sham. Red columns indicate upregulated (*3* on scaled expression), and blue columns indicate downregulated (*-3* on scaled expression). n = 4 per group (sham, *Serp1* WT and KD).

#### SUPPLEMENTARY FIGURE 4

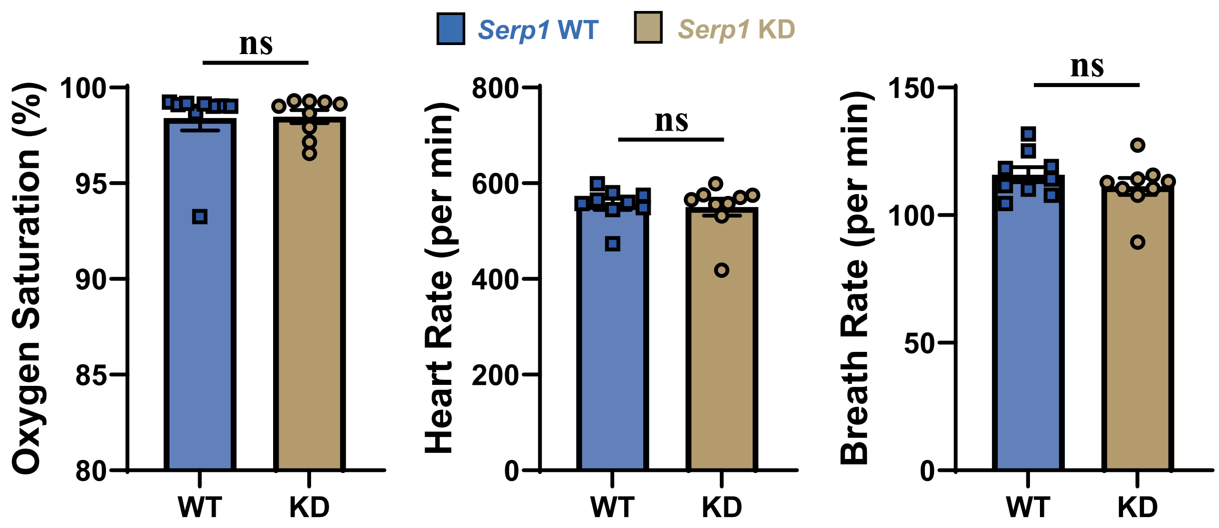


**Supp Fig. 4.** Analysis of mouse vital signs including oxygen saturation (%_, heart rate (per min), and breath rate (per min) in *Serp1* WT and KD mice 3d post stroke. Data analyzed with Mann-Whitney (nonparametric) t-test. ns p>0.05. Data expressed as the mean ± SEM.

#### SUPPLEMENTARY FIGURE 5

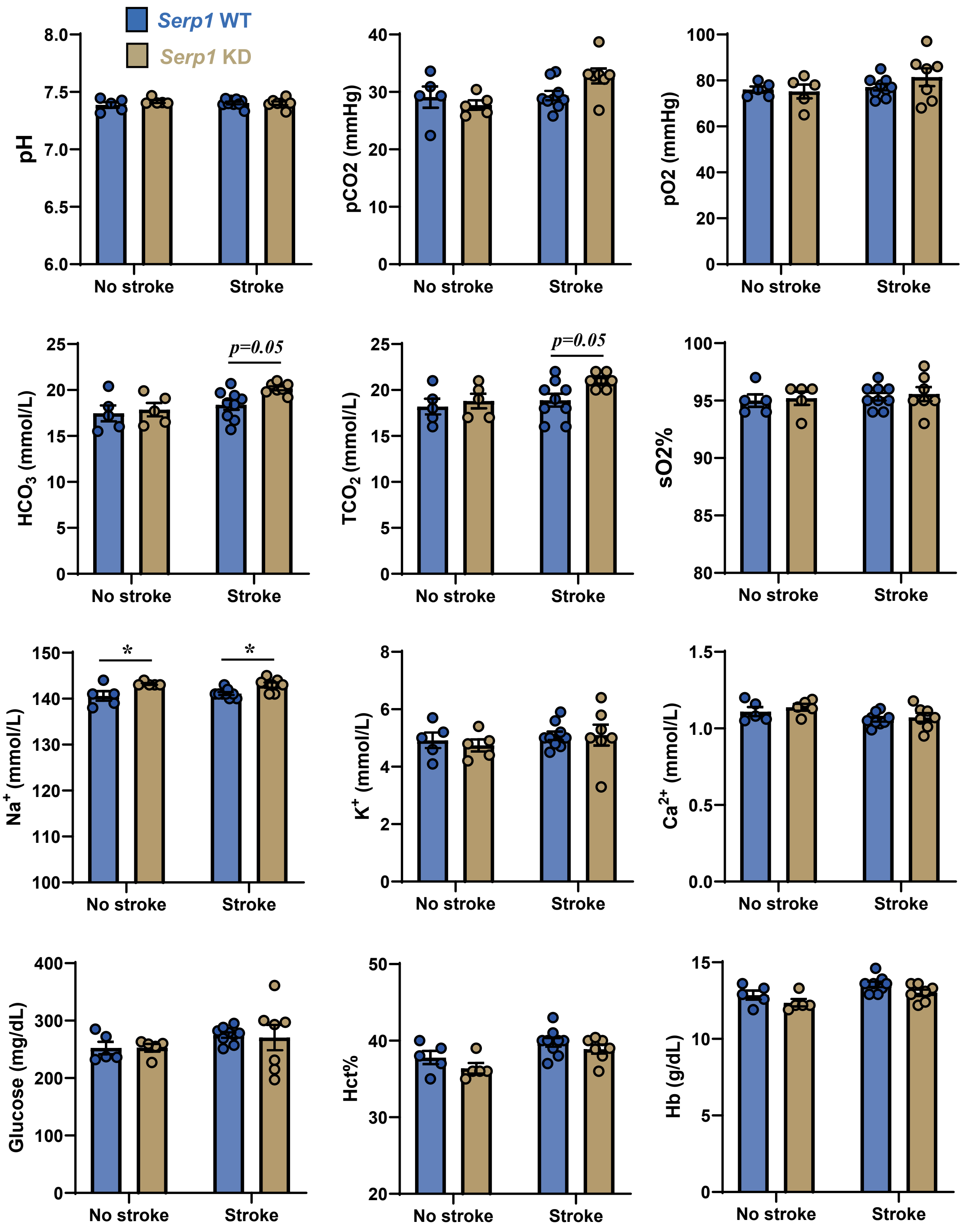


**Supp Fig. 5.** Analysis of blood chemistry in *Serp1* WT and KD mice in sham controls and 3d post stroke. Data analyzed with two-way ANOVA and Sidak's multiple comparisons test. *p<0.05. Data expressed as the mean ± SEM.

#### SUPPLEMENTARY FIGURE 6

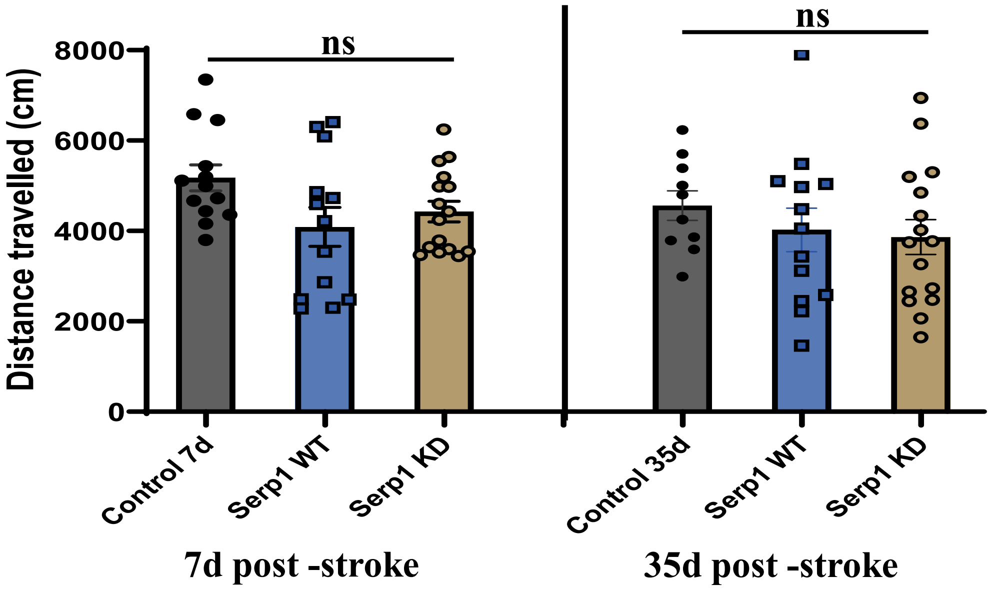


**Supp Fig. 6.** Open field behavior test between sham, Serp1 WT and KD mice performed at 7d- and 35d- post-stroke. Data analyzed with two-way ANOVA and Tukey's multiple comparisons test. ns $p>0.05$. Data expressed as the mean ± SEM.
